# A Mitochondrial Inside-Out Iron-Calcium Signal Reveals Drug Targets for Parkinson’s Disease

**DOI:** 10.1101/2022.10.30.513580

**Authors:** Vinita Bharat, Aarooran S. Durairaj, Roeland Vanhauwaert, Li Li, Colin M. Muir, Sujyoti Chandra, Chulhwan S. Kwak, Yann Le Guen, Pawan Nandakishore, Chung-Han Hsieh, Stefano E. Rensi, Russ B. Altman, Michael D. Greicius, Liang Feng, Xinnan Wang

## Abstract

Dysregulated iron or Ca^2+^ homeostasis has been reported in Parkinson’s disease (PD) models. Here we discover a connection between these two metals at the mitochondria. Elevation of iron levels causes inward mitochondrial Ca^2+^ overflow, through an interaction of Fe^2+^ with Mitochondrial Calcium Uniporter. In PD neurons, iron accumulation-triggered Ca^2+^ influx across the mitochondrial surface leads to spatially confined Ca^2+^ elevation at the outer mitochondrial membrane, which is subsequently sensed by Miro1, a Ca^2+^-binding protein. A Miro1 blood test distinguishes PD patients from controls and responds to drug treatment. Miro1-based drug screens in PD cells discover FDA-approved T-type Ca^2+^-channel blockers. Human genetic analysis reveals enrichment of rare variants in T-type Ca^2+^-channel subtypes associated with PD status. Our results identify a molecular mechanism in PD pathophysiology, and drug targets and candidates coupled with a convenient stratification method.

## Introduction

Parkinson’s disease (PD) is a leading cause of disability. The dopamine (DA)-producing neurons in the substantia nigra are the first to die in PD patients. A bottleneck that hinders our ability to effectively detect and treat PD may be the presence of highly heterogenous genetic backgrounds and risk factors among different patients. Functional studies have pointed to multiple “cellular risk elements” for PD, such as mitochondrial damage, lysosomal dysfunction, immune system activation, neuronal calcium mishandling, and iron accumulation ^1–10^. These distinct genetic and cellular risk factors may confer individual heterogeneity in disease onset, but also suggest that there are networks and pathways linking these “hubs” in disease pathogenesis. Identifying their connections could be crucial for finding a cure for PD.

Mitochondria are the center of cellular metabolism and communication. Ions such as Ca^2+^ and iron, are not only essential for diverse mitochondrial functions but can be stored inside the mitochondria to maintain cellular ionic homeostasis. Ion channels and transporters in the plasma and mitochondrial membranes coordinate for ion uptake, transport, and storage. Ca^2+^ enters the cell via voltage- or ligand-gated Ca^2+^ channels across the cell surface. Inside the cell, Ca^2+^ can be taken up by mitochondria through the outer mitochondrial membrane (OMM) channel, VDAC, and the inner mitochondrial membrane (IMM) channel, Mitochondrial Calcium Uniporter (MCU) ^11^, and extruded into the cytosol through the IMM transporter, NCLX ^12^. MCU is a multimeric holocomplex consisting of additional regulatory subunits, such as essential MCU regulator (EMRE), mitochondrial calcium uptake 1 (MICU1), MICU2, and MCUb ^13–15^, and can be substituted ^16^. For iron transport, iron bound carrier proteins such as transferrin deliver iron into the cell through endocytosis and intracellular iron can be taken up by mitochondria via several IMM transporters ^17^. Although individually, dysregulation of iron or Ca^2+^ homeostasis has been reported in PD models, their mechanistic link and contribution to disease susceptibility remain elusive. In this work, we harness the power of combining human genetics, cellular and in vivo models, and patient’s tissues, and identify a mitochondrial inside-out Fe^2+^-Ca^2+^-Miro1 axis in PD.

## Results

### Iron promotes the MCU complex assembly

The MCU’s Ca^2+^-import ability can be modulated by the elevation of cytosolic Ca^2+^ and intramitochondrial oxidants ^14,18^. We investigated whether the elevation of iron levels, a hallmark of PD ^9^, could also regulate MCU activity. It has been shown that following the stimulation of G-protein-coupled receptor (GPCR) agonist, the clearance of cytosolic Ca^2+^ is mediated by MCU ^18–20^. To investigate the relation of MCU and iron, we treated HEK293T cells with Fe^2+^, stimulated these cells the next day with the GPCR agonist, thrombin, and measured cytosolic and mitochondrial Ca^2+^ levels with live Calcium Green and Rhod-2 staining, respectively. We found that thrombin triggered intracellular Ca^2+^ mobilization and elevation, which was comparable between Fe^2+^-treated and mock control cells (Figure 1A-B). However, mitochondria in Fe^2+^-treated cells sustained a significantly larger Ca^2+^ elevation after thrombin stimulation, as compared to mock control (Figure 1C-D). These results indicate that iron promotes the mitochondrial Ca^2+^-import ability.

**Figure 1.**
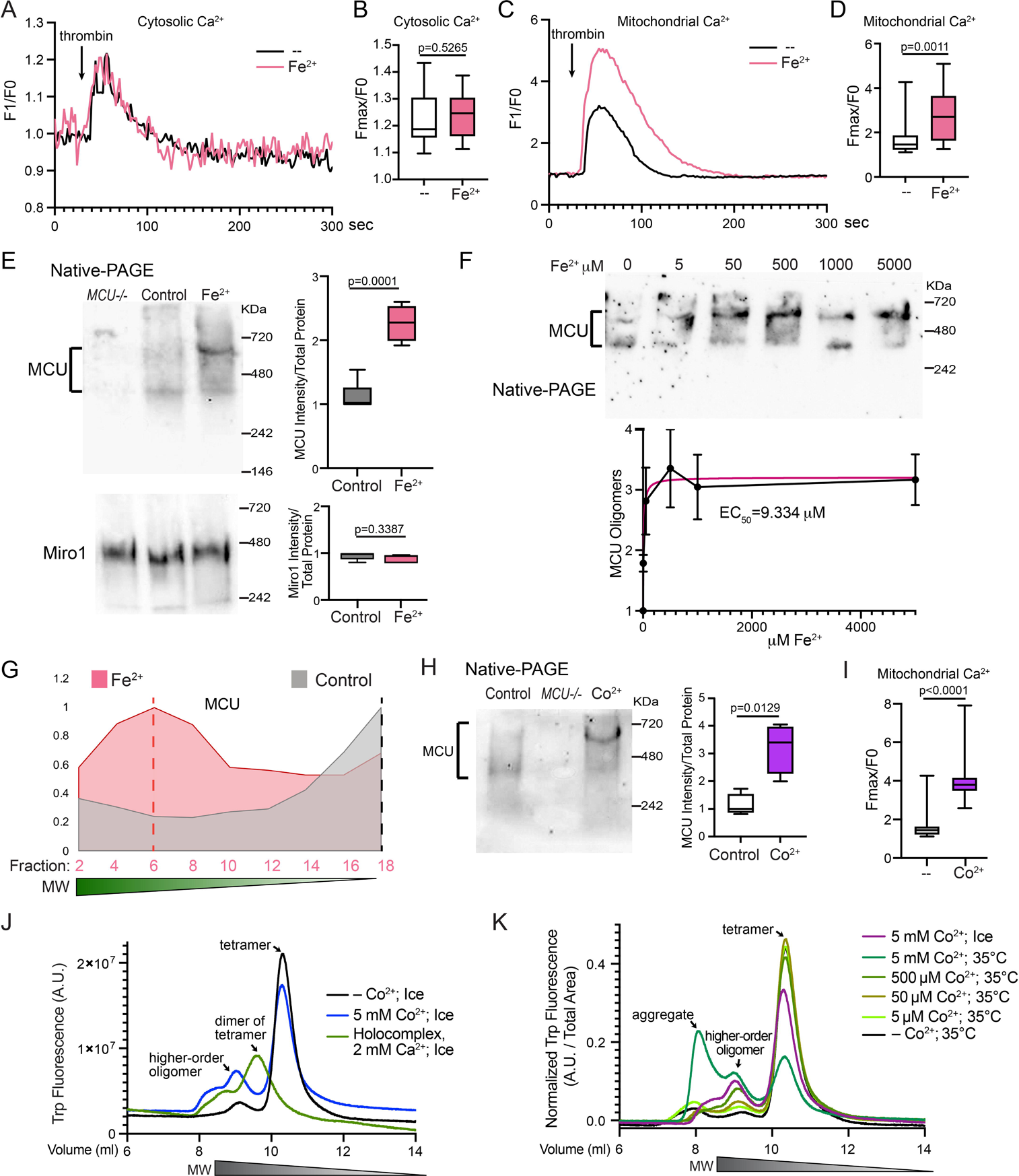
Iron Promotes MCU Oligomerization. (A-D) HEK cells were treated with 500 μM Fe^2+^ for 22 hours, stimulated with thrombin, and mitochondrial (Rhod-2) and cytosolic Ca^2+^ levels (Calcium Green) were measured. (A, C) Representative Ca^2+^ traces. (B, D) Quantification of the peak fluorescent intensity (Fmax) normalized to the baseline (F0) within each cell. n=20 cells from 4 independent coverslips. (E) HEK cells were treated with 5 mM Fe^2+^ for 22 hours, run in Native-PAGE, and blotted. Right: Qualification of the band intensity normalized to the total protein amount measured by a BCA assay. n=5. (F) Similar to (E), but with a range of Fe^2+^ concentrations added to the media. n=4. Data point shows Mean±S.E.M. (G) Elution profiles of MCU from SEC samples. (H) HEK cells were treated with 500 μM Co^2+^ for 22 hours and run in Native-PAGE. n=4. (I) HEK cells treated with 500 μM Co^2+^ for 22 hours were stimulated with thrombin and mitochondrial Ca^2+^ levels (Rhod-2) were measured. n=20 cells from 4 independent coverslips. (B, D, E, H, I) Two-tailed Welch’s T Test. (J-K) Fluorescence-detection SEC profiles. See also Figure S1.

Because MCU oligomerization affects its localization and potentially its Ca^2+^-uptake ability ^14,18^, we next explored whether iron altered the assembly of the MCU complex. Native-PAGE is a sensitive method to determine the overall form and amount of a multimeric native protein. The human MCU oligomer bands from HEK cells migrated between 400-700 KDa in Native-PAGE (Figure 1E) ^11,21,22^. Importantly, we found that Fe^2+^ treatment for 22 hours resulted in an increase in the total intensity of the MCU oligomer bands and appeared to shift some bands upward (Figure 1E), indicating the formation of higher-order oligomers. Miro1, an OMM protein, also oligomerized and migrated as a single band around 480 KDa in Native-PAGE, which was not significantly affected by Fe^2+^ treatment (Figure 1E). The EC_50_ of Fe^2+^ added to the media to cause the MCU response was 9.334 μM (Figure 1F). We confirmed that HEK cells were morphologically healthy under all concentrations and no significant cell death was observed (Figure S1A). To corroborate the Native-PAGE result, we performed an alternative method to differentiate protein complexes based on their molecular weight (MW)–size exclusion chromatography (SEC). Protein complexes with higher MW are eluted faster than those with lower MW. Detecting MCU from cell lysates using SEC has been successfully shown ^15,18,21^. We found that Fe^2+^ treatment shifted the MCU elution peak to the earlier fractions of higher-order oligomers compared to mock control (Figure 1G, S1B; anti-MCU was validated in Figure S1C). As a positive control for this method, we treated cells with H_2_O_2_ for 10 minutes and found the shift of the MCU elution peak to the earlier fractions (Figure S1D), consistent with previous studies also using SEC ^18^. In contrast, the elution pattern of Miro1 was largely unaltered by any of these treatments (Figure S1E). These data indicate that iron shifts the MCU complexes to higher-order oligomers and enlarges the total number of these complexes.

To dissect the role of reactive oxygen species (ROS) ^18^ in contributing to the Fe^2+^-triggered MCU function as observed above, we treated HEK cells with an antioxidant, Vitamin C (VitC) ^23^, which has no iron-chelating ability ^24^, or an iron chelator, deferiprone (DFP) ^25^, together with Fe^2+^, and measured MCU oligomerization and mitochondrial Ca^2+^ uptake. DFP, but not VitC, lowered the MCU oligomer band intensity and mitochondrial Ca^2+^ import in the presence of Fe^2+^ (Figure S1F-G), indicating that ROS might have a nominal effect on Fe^2+^-mediated MCU regulation, at least in this experimental setting.

We attempted to reconstitute the phenomenon of the Fe^2+^-triggered increase of MCU oligomerization in vitro. However, a fast speed of Fe^2+^ oxidation caused precipitation of purified MCU protein. To circumvent this problem, we used an Fe^2+^ mimic, Co^2+^ ion ^26^. The data of Co^2+^ might inform us about properties of Fe^2+^. We confirmed that Co^2+^ behaved similarly as Fe^2+^ in our functional assays in HEK cells: Co^2+^ treatment increased MCU oligomerization detected by Native-PAGE (Figure 1H) and enhanced the mitochondrial Ca^2+^-uptake ability following thrombin application (Figure 1I), just like Fe^2+^ (Figure 1C-E). We then conducted fluorescence-detection SEC using purified human MCU protein ^14^, and found that Co^2+^ caused the formation of higher-order oligomers of MCU (Figure 1J). As a positive control for this approach, we reproduced the elution peak of the dimer of tetramer of the MCU holocomplex caused by Ca^2+^ addition in vitro (Figure 1J) ^14^. These results demonstrate that the Fe^2+^ mimic, Co^2+^, increased MCU oligomerization in vitro. Collectively, Fe^2+^ and its mimic Co^2+^ promote the formation of MCU higher-order oligomers and enhance the mitochondrial Ca^2+^-uptake activity.

### Fe^2+^ binds to the MCU complex to promote its oligomerization and channel activity

Our in vitro result in Figure 1J suggested that Co^2+^ bound to MCU to cause the formation of higher-order oligomers. We further confirmed that Co^2+^ altered MCU oligomerization and thermostability in a dose-dependent manner (Figure 1K).

We hypothesized that Fe^2+^ also bound to MCU to increase its oligomerization. To explore this possibility, we immunoprecipitated (IP) endogenous MCU from HEK cells and detected the iron concentrations in the IP samples. We found significantly more Fe^2+^ ions pulled down with MCU when HEK cells were treated with Fe^2+^ (Figure 2A). We next examined whether MCU directly bound to Fe^2+^ by an in vitro binding assay. To minimize protein precipitation caused by Fe^2+^ oxidation, we performed an alternative to the thermal shift-SEC experiment with Co^2+^ shown in Figure 1K, which could take hours to complete. We applied Fe^2+^ to the purified MCU complex (MCU, MICU1, MICU2, ERME) ^14^ for about 5 minutes and after wash immediately measured iron levels in the complex by inductively coupled plasma mass spectrometry (ICP-MS). We found that the MCU complex bound to Fe^2+^ in vitro with a Kd of 449.3 μM (Figure 2B).

**Figure 2.**
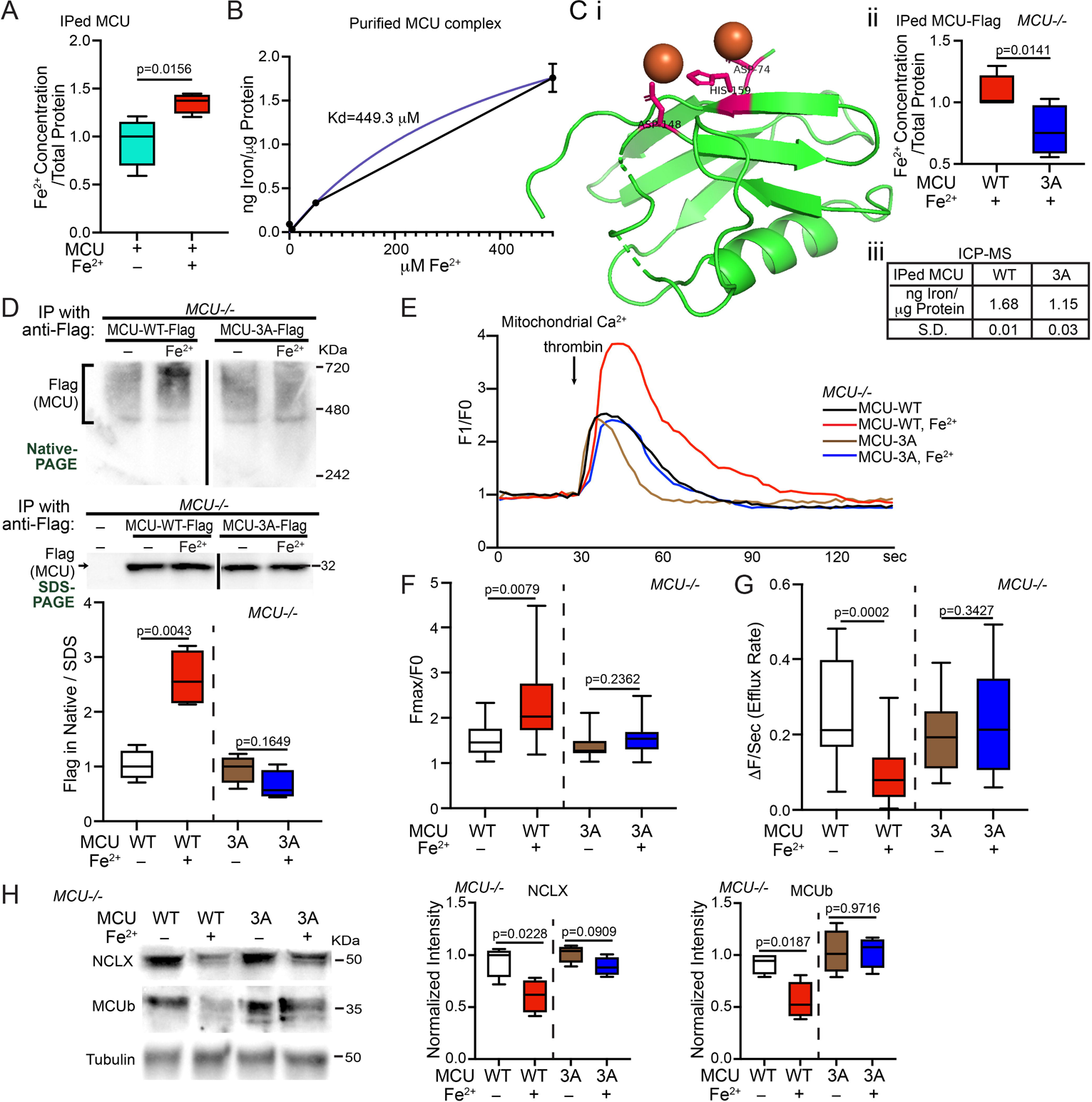
MCU Binds to Fe^2+^. (A) HEK cells were treated with or without 500 μM Fe^2+^ for 21 hours, then IPed with anti-MCU, and Fe^2+^ concentrations in the IP samples were detected (normalized to the total protein amount). Two-tailed paired T Test. n=4. (B) Different concentrations of Fe^2+^ was added to the purified MCU complex in vitro and after wash the total iron level in the complex was detected by ICP-MS. Mean±S.E.M. n=3. (Ci) Structural visualization of the matrix domain of MCU (green) bound to Fe^2+^ (orange) via these sites (pink) generated by PyMol. (Cii) *MCU-/-* HEK cells transfected as indicated were treated with 500 μM Fe^2+^ for 20 hours, then IPed with anti-Flag, and Fe^2+^ concentrations in the IP samples were detected. Two-tailed paired T Test. n=4. (Ciii) ICP-MS measurement of the total iron level in IPed MCU-Flag from cells as in (Cii), normalized to the total protein amount. n=3. (D) Representative blots of IP with anti-Flag using cell lysates as indicated, run in Native- or SDS-PAGE. n=4. (E) *MCU-/-* HEK cells transfected as indicated and treated with or without 500 μM Fe^2+^ for 22 hours were stimulated with thrombin and mitochondrial Ca^2+^ levels (Rhod-2) were measured. Representative Ca^2+^ traces. (F-G) Based on traces like in (E), the peak fluorescent intensity normalized to baseline (F) or efflux rate (G) is quantified. n=17 cells from 4 independent coverslips. (H) Cells as in (D-E) were lysed and blotted. Normalized to tubulin. n=4. (D-H) Two-tailed Welch’s T Test. Relative to “MCU-WT” or “MCU-3A” without treatment.

We next searched for amino acid residues in the matrix domain of MCU (PDB: 5KUE) predicted to bind to Fe^2+^ using an in-silico program (http://bioinfo.cmu.edu.tw/MIB/) ^27,28^, and found 3 amino acids: 74D, 148D, and 159H (Figure 2Ci). The latter 2 residues were also predicted to bind to Co^2+^. We mutated these 3 sites to Alanine (named “MCU-3A”). If MCU bound to Fe^2+^ via some of these sites, MCU-3A should reduce their binding. Indeed, we detected significantly fewer Fe^2+^ in the immunoprecipitate of Flag-tagged MCU-3A, as compared to MCU-WT, expressed from HEK cells without endogenous MCU (*MCU-/-*) treated with Fe^2+^ (Figure 2Cii). We confirmed this result by ICP-MS (Figure 2Ciii). Thus, MCU may bind to Fe^2+^ through some of these sites.

To determine whether blocking MCU’s binding to Fe^2+^ was sufficient to eliminate the Fe^2+^-triggered oligomerization of MCU, we expressed MCU-WT or MCU-3A in *MCU-/-* HEK cells to prevent the interference of endogenous MCU, treated these cells with Fe^2+^, and ran IPed MCU in Native-PAGE. As expected, MCU-3A abolished MCU’s response to Fe^2+^ treatment: the MCU oligomer band intensity was no longer increased (Figure 2D). We then live imaged mitochondrial Ca^2+^ dynamics, as described in Figure 1, in these cells. We consistently observed a larger mitochondrial Ca^2+^ elevation following thrombin stimulation in MCU-WT-transfected HEK cells treated with Fe^2+^ as compared to no Fe^2+^ treatment, and MCU-3A blunted the peak increase (Figure 2E-F). Of note, MCU-3A-expressing cells maintained normal functions of MCU (Figure 2D-H), suggesting no negative impact on cells by these mutations. In addition, ROS similarly promoted the oligomerization of MCU-WT and MCU-3A (Figure S1H), consistent with that the cysteine residues for oxidation ^18^ remain intact in both proteins. Thus, ROS might have a negligible role in the difference between MCU-WT and 3A triggered by Fe^2+^ seen in Figure 2D-H. Altogether, our results show that Fe^2+^ promotes MCU oligomerization and its channel activity through binding to MCU.

### Iron reduced NCLX and MCUb protein levels in a manner dependent on MCU channel activity

We next examined additional membrane proteins that may assist MCU function using HEK cells. By detecting total protein levels using Western blotting, we found a reduction in only MCUb and NCLX but not the other 12 proteins examined including MCU monomers, when cells were treated with Fe^2+^ for 20-22 hours (Figure S2), demonstrating selective protein responses without widespread toxicity at this time point (Figure S1A). MCUb is an inhibitor of MCU ^15^, and NCLX is an IMM exchanger believed for mitochondrial Ca^2^ extrusion ^12,29^. The reduction of both proteins could contribute to the phenotype of mitochondrial Ca^2+^ overload upon Fe^2+^ treatment observed earlier (Figure 1C-D).

We explored the possibility that the reduction of NCLX and MCUb was secondary to MCU activity and resultant Ca^2+^ overload. We took advantage of the *MCU-3A*-expressing *MCU-/-* cells which eliminated MCU’s Fe^2+^-binding and Fe^2+^-elicited channel activity (Figure 2). By measuring the Ca^2+^ extrusion rate we found that Fe^2+^ treatment slowed the efflux rate in *MCU-WT*-expressing *MCU-/-* cells, which was prevented by MCU-3A (Figure 2G). This result suggests that the Fe^2+^-triggered efflux delay may depend on Ca^2+^ overload caused by increased MCU activity. We then immunoblotted NCLX and MCUb protein from these cells. We found again the reduction of NCLX and MCUb in *MCU-WT*-expressing *MCU-/-* cells upon Fe^2+^ treatment and their protein levels were restored in *MCU-3A* expressing cells (Figure 2H). Hence, these findings indicate that mitochondrial Ca^2+^ overload upon Fe^2+^ elevation is the primary reason for the reduction of NCLX and MCUb levels.

### PD postmortem brains and neurons mirror Fe^2+^-treated HEK cells

Our data so far have shown that artificially increasing iron levels causes mitochondrial Ca^2+^ accumulation by promoting MCU activity. We examined this mechanism in pathophysiological PD models without the need of adding exogenous iron to the media. We first investigated postmortem brains of people with sporadic PD without a family history (Table S1). We homogenized the frontal cortex, where iron elevation was observed in PD patients before measured by ICP-MS ^30^, and ran the brain lysate in Native- or SDS-PAGE. Consistent with previous studies, MCU, NCLX, and MCUb were detected in the human brain ^8,31^. We found that the PD group (7 patients) clustered and separated from the healthy control group (5), with higher intensity of the MCU oligomer bands in Native-PAGE and lower intensity of both the NCLX and MCUb bands in SDS-PAGE (Figure 3A, S3A), similar to the observations in HEK cells treated with Fe^2+^ (Figure 1, S2). This unique clustering was not observed in the Alzheimer’s disease (AD) group (5 patients) (Figure S3A). The total iron level in each brain fraction, measured by ICP-MS, was positively correlated with the corresponding band intensity of MCU oligomers in Native-PAGE (Figure 3B).

**Figure 3.**
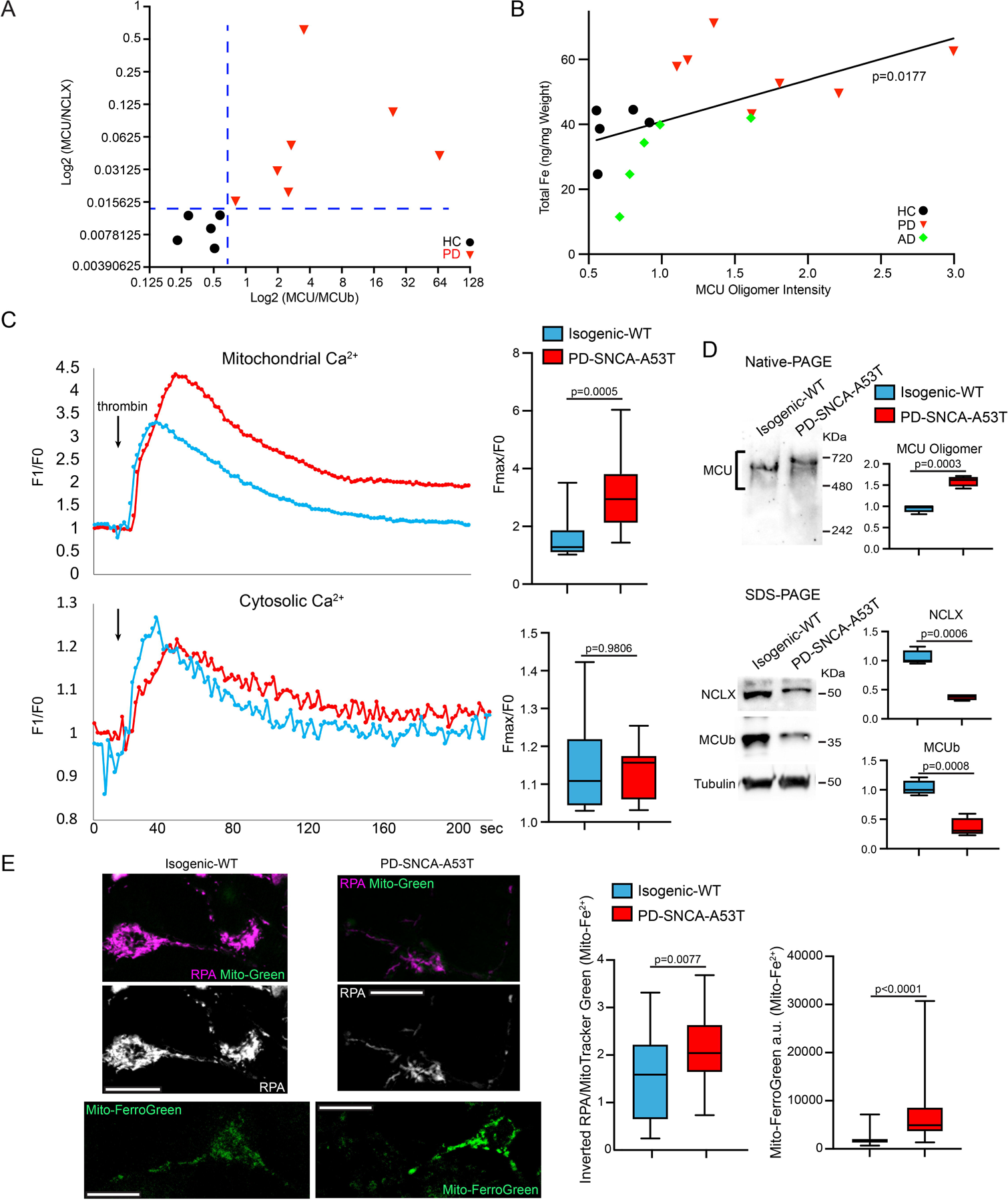
PD Models Mirror Fe^2+^-Treated HEK Cells. (A) Postmortem brains were run in Native- or SDS-PAGE and blotted. Blots and details are in Figure S3A. HC: healthy control. (B) The total iron level (y axis) in each brain fraction was measured by ICP-MS (normalized to wet weight, average of 3 repeats), and plotted with the band intensity of MCU oligomers in Native-PAGE (x axis) from the same brain fraction. Linear fit. (C) iPSC-derived neurons were stimulated with thrombin and mitochondrial (Rhod-2) and cytosolic Ca^2+^ levels (Calcium Green) were measured in the somas. Left: Representative Ca^2+^ traces. n=15 cell bodies from 3 independent coverslips. (D) iPSC-derived neurons were lysed and blotted as indicated. Normalized to the total protein amount. n=4. (E) Left: Confocal single section images of live neurons as in (C-D) stained with RPA (quenched by Fe^2+^) and MitoTracker Green (Mito-Green), or Mito-FerroGreen (stains Fe^2+^). Scale bars: 10 μm. Middle: Quantification of the RPA fluorescent intensity in each soma from inverted images normalized to MitoTracker Green, a membrane-potential-independent dye whose intensity was similar between control and PD neurons (P=0.2524). n=26 (WT) and 28 (PD) cell bodies from 3 coverslips. Right: Quantification of the Mito-FerroGreen fluorescent intensity in each soma subtracting background. n=55 cell bodies from 3 coverslips. Two-tailed Welch’s T Test. See also Figure S3.

We next employed induced pluripotent stem cells (iPSCs) from one familial patient (PD-SNCA-A53T) with the A53T mutation in *SNCA* (encodes α-synuclein) and its isogenic wild-type control (isogenic-WT) ^32–35^. We differentiated iPSCs to neurons expressing tyrosine hydroxylase (TH), the rate-limiting enzyme for DA synthesis as previously described ^32–35^. These patient-derived neurons display increased expression of endogenous α-synuclein ^33^. We stimulated these neurons with thrombin and measured cytosolic and mitochondrial Ca^2+^ levels as in Figure 1. We found very similar Ca^2+^ dynamic pattern in PD neurons as in HEK cells treated with Fe^2+^: PD neurons suffered from more mitochondrial Ca^2+^ accumulation after thrombin stimulation, as compared to isogenic control (Figure 3C). We next ran the neuronal lysates in Native- and SDS-PAGE. We again observed an upward shift of the MCU oligomer bands and an increase in the total MCU band intensity in Native-PAGE indicating the formation of higher-order oligomers, and a reduction of the intensity of NCLX and MCUb bands in SDS-PAGE in PD neurons compared to control (Figure 3D). We also discovered a significant increase in the mitochondrial Fe^2+^ levels detected by RPA and Mito-FerroGreen staining in PD neurons (Figure 3E). The phenotypes of mitochondrial Fe^2+^ and Ca^2+^ were observed in older neurons as well (day 40, Figure S3B). Collectively, our results demonstrate a positive correlation of iron levels with MCU oligomerization and its channel activity in PD patient-derived neurons and postmortem brains.

### Iron functions upstream of calcium to mediate phenotypes of PD neurons

Our findings suggested the possibility that the elevation of iron levels in PD caused mitochondrial Ca^2+^ overload by the binding of MCU with Fe^2+^ (Figures 1-3). To test this hypothesis, we reduced iron levels in PD neurons described in Figure 3 with DFP and found that DFP significantly lowered mitochondrial Ca^2+^ accumulation following thrombin stimulation (Figure 4A), demonstrating a causal relation between iron elevation and Ca^2+^ overload in the mitochondria. These PD neurons are more vulnerable to mitochondrial stressors. The treatment of the complex III inhibitor, Antimycin A, at 10 μM for 6 hours caused acute neuronal cell death leading to the increase of terminal deoxynucleotidyl transferase dUTP nick end labeling (TUNEL) signals in neurons derived from this PD patient, but not in isogenic control neurons (Figure S3C) ^33–35^. We found that pretreatment with DFP prevented cell death (Figure S3C). We validated these findings in neurons differentiated from additional iPSC lines, including those from two sporadic patients (PD-S1, PD-S2; no family history) and two healthy controls (HC-1, HC-2) ^32–34^, and an isogenic pair containing a mutant line (Jax-SNCA-A53T) where *SNCA-A53T* was made in iPSCs of a healthy donor and an isogenic control (Jax-Rev) by reverting *SNCA-A53T* back to wild type (Method). We consistently observed key features shown earlier: mitochondrial Fe^2+^ and stimulated-Ca^2+^ levels were elevated (Figure S3D-E), and DFP prevented mitochondrial Ca^2+^ overload (Figure S3D) and stressor-induced cell death (Figure S3C) in these PD-related lines. In order to cross-validate the neuroprotection of DFP in vivo without the need of adding a stressor, we fed DFP to a fly model of PD, which expressed the pathogenic human α-synuclein protein with the A53T mutation (α-syn-A53T) in DA neurons driven by *TH-GAL4* ^33–35^. These flies exhibit PD-relevant phenotypes such as age-dependent locomotor decline and DA neuron loss ^33–35^. In the protocerebral posterior lateral 1 (PPL1) cluster of the fly brain, the DA neuron number typically ranges between 11-14, but this number is reduced in PD flies. We found that feeding PD flies with DFP restored the DA neuron number in aged fly brains and improved the climbing ability (Figure 4B-C). Collectively, our results show that iron functions upstream of calcium and mediates neurodegeneration in multiple PD models.

**Figure 4.**
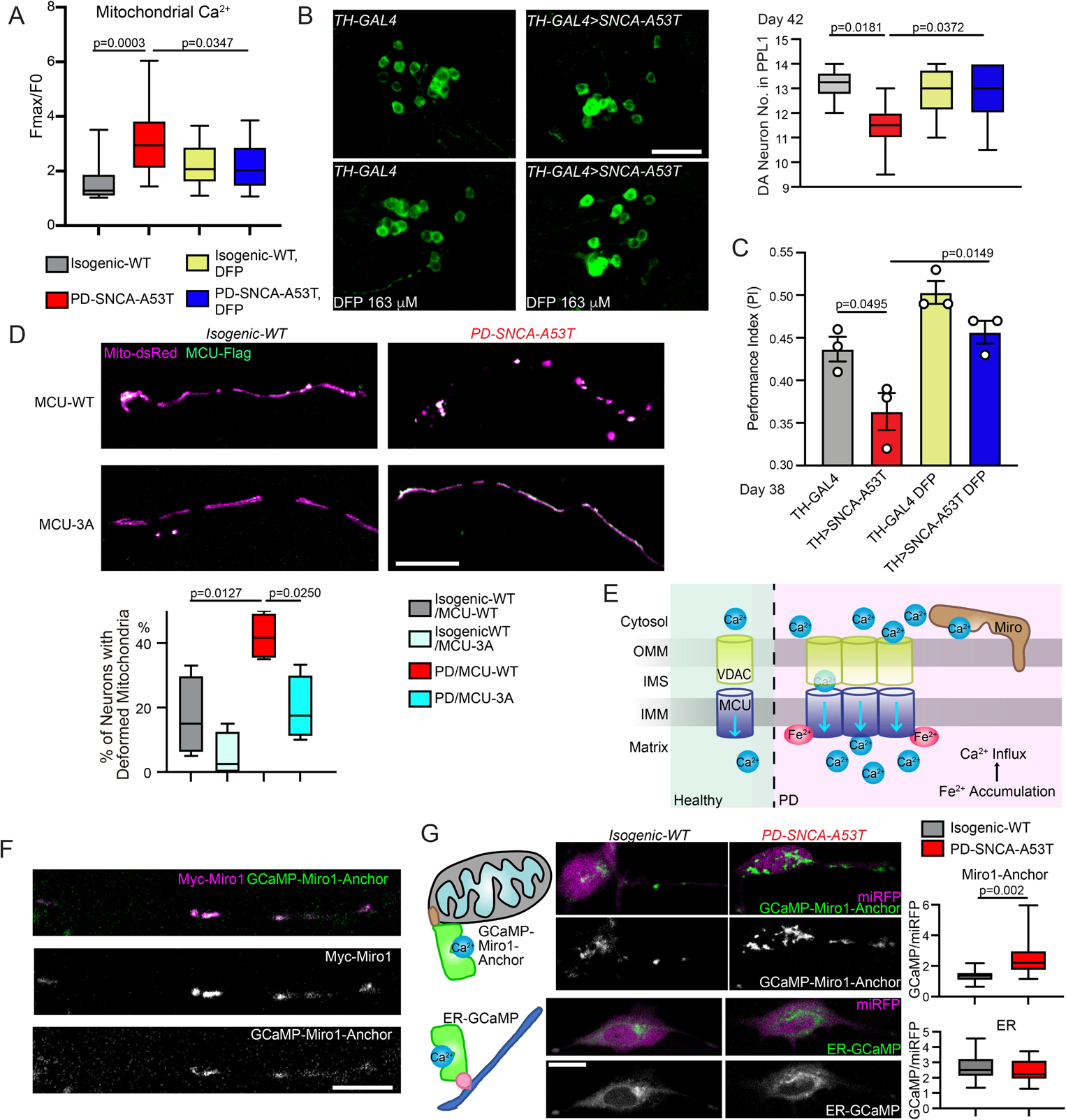
Iron Functions Upstream of Calcium in PD neurons. (A) Similar to Figure 3, iPSC-derived neurons, with or without treatment of 100 μM DFP for 24 hours, were stimulated with thrombin, and mitochondrial Ca^2+^ (Rhod-2) was measured. n=15 cell bodies from 3 independent coverslips. Control data without DFP treatment are the same as in Figure 3C. (B) The DA neuron number (average of both sides) was counted in the PPL1 cluster. Drug treatment was started from adulthood (day 1). Scale bar: 20 μm. n=6, 9, 8, 7 (from left to right) flies. (C) Drug treatment was started from embryogenesis. n=35, 33, 40, 34 flies (from left to right), 3 independent experiments. (D) Representative confocal images of neurites transfected as indicated. Scale bar: 10 μm. Quantification of the percentage of total neurons with deformed mitochondria (defined as a neuron without any mitochondrion in neurites >2.5 μm in length). n=4 coverslips. (A, D) One-Way Anova Post Hoc Tukey Test. (B-C) Two-Way Anova Post Hoc Tukey Test. (E) Schematic representation of iron triggered mitochondrial calcium influx in PD. (F) A *PD-SNCA-A53T* iPSC-derived neuron was transfected as indicated and immunostained with anti-Myc. Scale bar: 10 μm. (G) Confocal live images of neurons transfected as indicated. Scale bar: 10 μm. Quantification of the intensity of GCaMP6f normalized to miRFP670 within the same cell body. n=15 cell bodies from 3 coverslips. Two-tailed Welch’s T Test. See also Figure S3.

We next utilized the Fe^2+^-resistant MCU mutant, MCU-3A (Figure 2), to block extra Fe^2+^-binding in PD neurons. While expressing MCU-WT in PD neurons caused mitochondrial deformation including shortening, fragmenting, and swelling, probably due to substantial iron-mediated mitochondrial Ca^2+^ overload ^36^, MCU-3A completely prevented this phenotype (Figure 4D). As mentioned earlier, ROS should similarly oxidize both MCU-WT and 3A proteins ^18^ (Figure S1H) and impact those PD neurons. These results suggest that eliminating MCU’s binding to Fe^2+^ renders protection for mitochondria.

Together, our data from both HEK cells and PD models demonstrate a pathological mechanism whereby iron accumulation inside the mitochondria causes inward Ca^2+^ overflow via a direct binding of Fe^2+^ with MCU and the resultant augmentation of MCU activity (Figure 4E).

### Local OMM calcium dysregulation is sensed by Miro1 in Parkinson’s models

We reasoned that the enlarged Ca^2+^ ion influx through MCU in PD models could alter local Ca^2+^ dynamics and balance at the OMM (Figure 4E). This subdomain-scale Ca^2+^ change might be sensed by a nearby OMM protein. Miro1 is a Ca^2+^-binding protein on the OMM facing the cytosol, and in close proximity to mitochondrial and ER Ca^2+^ channels ^2,37–39^. Thus, Miro1 may directly encounter Ca^2+^ flux going through mitochondria and sense even spatially confined Ca^2+^ concentration changes. To explore this possibility, we swapped the cytosolic portion of Miro1 with the fluorescent Ca^2+^ sensor, GCaMP6f, while leaving Miro1’s C-terminal transmembrane and tail domain intact to allow proper OMM localization ^35^, named GCaMP-Miro1-Anchor. A cytosolic expressing miRFP670 gene was constructed in the same vector for background control. This Ca^2+^ sensor should detect Ca^2+^ ions at the subdomains of the OMM where Miro1 is located. We confirmed that GCaMP-Miro1-Anchor was colocalized with Miro1 in human neurons (Figure 4F). The intensity of GCaMP-Miro1-Anchor was increased showing elevated local Ca^2+^ concentrations in live PD neurons compared to isogenic control, while the intensity of a Ca^2+^ sensor located to the cytosolic face of the ER membrane was unchanged (Figure 4G). This increase in the GCaMP-Miro1-Anchor intensity was not caused by increased mitochondrial volume because the fluorescent intensity of MitoTracker Green, a membrane-potential-independent dye, was comparable between control and PD neurons (Figure 3E; Figure Legend). Importantly, inhibiting the MCU activity by RU360 ^40^ reduced OMM Ca^2+^ elevation detected by GCaMP-Miro1-Anchor in PD neurons (Figure S3F). This data suggests that enhanced mitochondrial Ca^2+^ influx could raise spatially restricted Ca^2+^ concentrations at the OMM and that Miro1 may serve as an ideal sensor for this change (Figure 4E).

We next sought a quantitative measurement of Miro1’s Ca^2+^ sensitivity at the OMM, which could help us develop specific molecular markers to distinguish PD patients with OMM Ca^2+^ elevation. We had previously found a Miro1 phenotype in PD models and patients’ cells. We explored whether this Miro1 phenotype was a result of Miro1’s Ca^2+^ sensing. In a healthy cell, upon mitochondrial depolarization Miro1 is quickly removed from the OMM and degraded in proteasomes, primed by PD-related proteins including PINK1, Parkin, and LRRK2, allowing the subsequent mitophagy ^32–34,41,42^. We had shown that Miro1 degradation upon mitochondrial depolarization was delayed in fibroblasts or neurons derived not only from familial PD patients with mutations in *Parkin* or *LRRK2* but also from sporadic patients without known mutations, consequently slowing mitophagy and increasing neuronal sensitivity to stressors ^32–34^. The mechanism underlying this unifying Miro1 phenotype in both familial and sporadic PD patients remained undefined. To explore whether this phenotype resulted from Miro1’s Ca^2+^ binding, we made GFP-tagged human Miro1 protein in both the wild-type form (Miro1-WT) and in a mutant form where two point mutations were introduced in the two EF-hands of Miro1 (Miro1-KK) to block Ca^2+^ binding ^43^. We expressed GFP-tagged Miro1 (WT or KK) and Mito-dsRed that labeled mitochondria in iPSC-derived neurons from the PD patient and isogenic control, described earlier. We had previously discovered that following acute Antimycin A treatment that depolarized the membrane potential and triggered mitophagy, Miro1 and mitochondria were sequentially degraded in wild-type neurons ^32–34^. We observed the same mitochondrial events in isogenic control axons transfected with GFP-Miro1-WT by live imaging (Figure 5A-C). In contrast, in PD neuron axons transfected with GFP-Miro1-WT, the degradation rates of both Miro1 and damaged mitochondria upon Antimycin A treatment were slowed (Figure 5A-C), consistent with our previous studies ^33,34^. Notably, GFP-Miro1-KK significantly rescued these phenotypes in PD axons: it expedited the degradation rates to the control level (Figure 5A-C). We validated the result of Miro1 degradation using an enzyme-linked immunosorbent assay (ELISA) of GFP (Figure 5D). These data suggest that delayed Miro1 and damaged mitochondrial degradation rely on Miro1’s Ca^2+^ binding in PD neurons.

**Figure 5.**
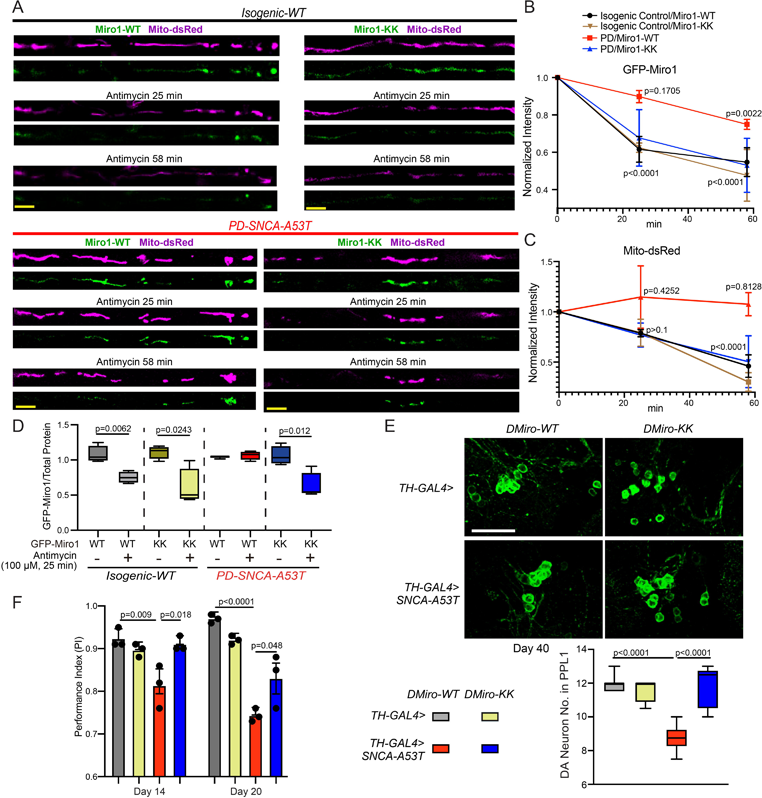
Miro Senses Calcium to Mediate Several PD Relevant Phenotypes. (A) Representative still images from live Mito-dsRed and GFP-Miro1 recordings in axons of indicated genotypes, following 100 μM Antimycin A treatment. Scale bar: 10 μm. (B-C) Degradation rate profiles of GFP-Miro1 (B) or Mito-dsRed (C) normalized to the same axonal region at 0 min. n=5 axons (one axon per coverslip). Comparison with “0 min”. One-Way Anova Post Hoc Dunnett’s Test. (D) iPSC-derived neurons were transfected as indicated and GFP-Miro1 was detected by an ELISA normalized to the total protein amount. n=4. Two-tailed Welch’s T Test within each condition. (E) The DA neuron number. Scale bar: 20 μm. n=7, 4, 6, 5 (from left to right). (F) n (from left to right)=49, 47, 39, 47 (day 14); 48, 45, 37, 44 (day 20); 3 independent experiments. (E-F) Two-Way Anova Post Hoc Tukey Test. See also Figure S3.

To confirm the Miro-Ca^2+^ relation in vivo, we generated transgenic flies carrying T7-tagged fly Miro (DMiro)-WT or DMiro-KK. DMiro is an ortholog of human Miro1 with high sequence similarity. Both DMiro-WT and DMiro-KK were expressed at comparable levels when the transgenes were driven by the ubiquitous driver *Actin-GAL4* (Figure S3G). We next crossed these transgenic flies to a fly PD model described earlier that expressed human α-syn-A53T in DA neurons driven by *TH-GAL4*. We found that DMiro-KK significantly rescued the phenotypes of age-dependent DA neuron loss and locomotor decline, compared to DMiro-WT (Figure 5E-F). This result shows that blocking DMiro’s Ca^2+^-binding is neuroprotective in PD flies. Altogether, we have provided evidence that Miro1 or DMiro senses Ca^2+^ to mediate several phenotypes in human neuron and fly models of PD.

### A high-content Miro1 screening assay identifies a network of Ca^2+^-related drug hits for PD

To further support that Miro1 was downstream of Ca^2+^ dysregulation in PD, we performed high throughput (HTP) drug screens in fibroblasts from a sporadic PD patient using the Miro1 phenotype (delayed degradation upon depolarization) as a readout. If Miro1 sensed local Ca^2+^ elevation to become more stable on damaged mitochondria (Figure 4G, 5A-C), we expected to see Ca^2+^-related drug hits to show up to promote Miro1 degradation. To this end, we established a sensitive immunocytochemistry (ICC)-based assay that was suitable for HTP screening (Figure 6A, S4, S5, S6A, Table S2, Method). We performed the primary screens at the Stanford High-Throughput Bioscience Center (HTBC) using 3 drug libraries containing many compounds that have well-defined roles and targets and show efficacy to treat certain human diseases (Figure S4). Drug hits from primary screens were then retested in our own laboratory using fresh compounds at the highest screening concentration with at least 4 biological replicates and our confocal microscope (Figure S5). In total, we discovered 15 compounds in two independent experimental settings that reduced Miro1 protein levels following mitochondrial depolarization in PD fibroblasts (Figure S5C, Table S2).

**Figure 6.**
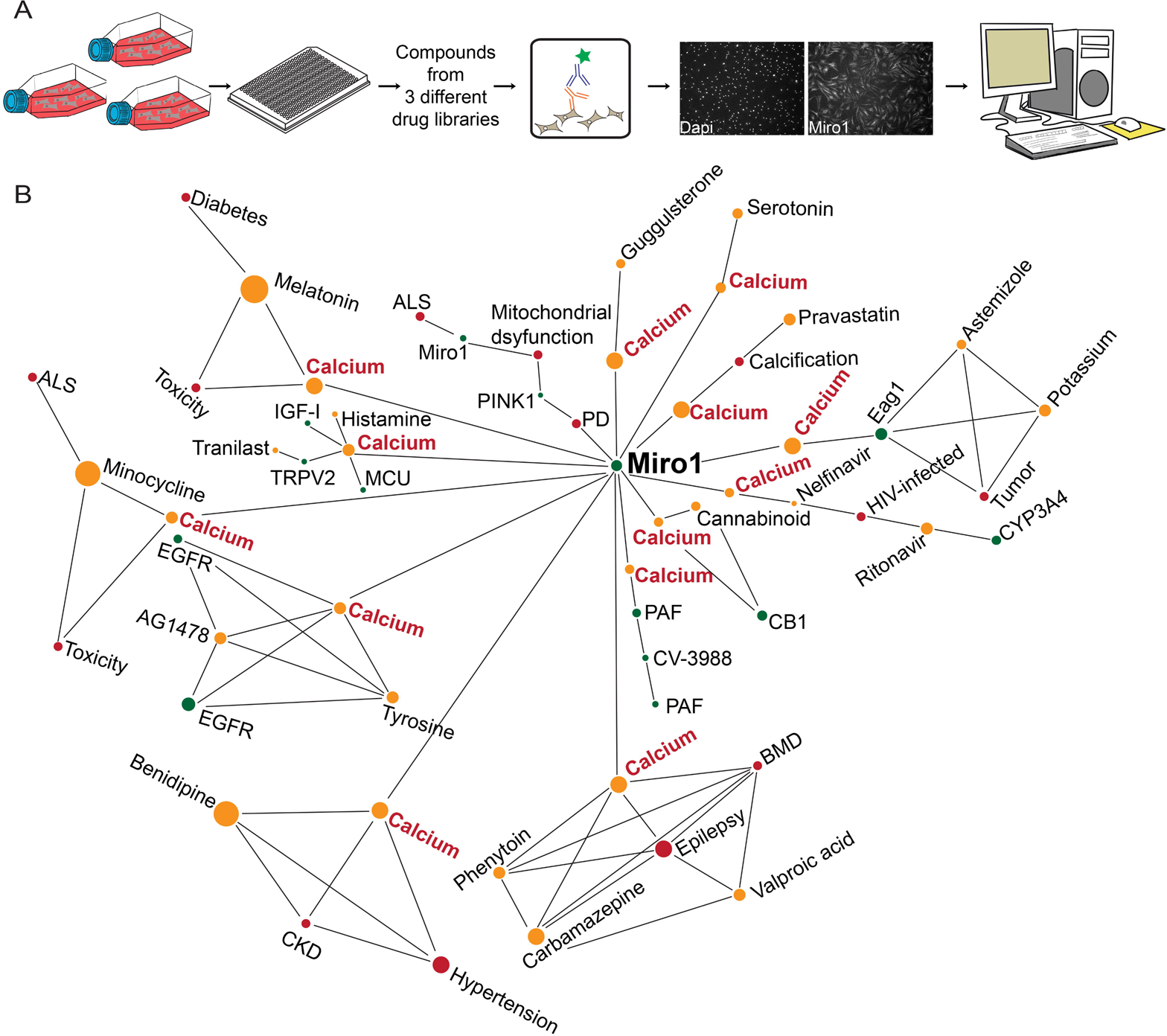
HTP Screens Identify Ca^2+^-Related Drug Hits for PD. (A) Schematic representation of a custom-designed drug screen for Miro1 in PD fibroblasts. (B) Pathway analysis identifies Ca^2+^ as a shared factor in the primary hit-Miro1 network. See also Figure S4-5.

Next, we performed a pathway analysis using a knowledge graph-based tool to reveal the potential cellular pathways connecting Miro1 to each hit compound. Strikingly, we discovered intracellular Ca^2+^ as a primary shared factor in the hit drug-Miro1 network (Figure 6B, Table S3). Two drugs, Benidipine and Tranilast, could directly inhibit plasma membrane Ca^2+^ channels. Benidipine is a blocker of voltage-gated Ca^2+^ channels (L-, N-, T-type), and Tranilast has been proposed to inhibit ligand-gated Ca^2+^ channels (TRPV2) ^44^. Therefore, our results confirm the importance of the Ca^2+^ pathway for mediating the Miro1 phenotype in PD.

### Benidipine rescues Parkinson’s-relevant phenotypes in multiple models of PD

We further validated the top hit from our screens, Benidipine, a pan-Ca^2+^-channel blocker. Using the same ICC method as in Figure S5, we found that Benidipine reduced Miro1 in a dose-dependent manner upon depolarization in PD fibroblasts (Figure S6B). To exclude the possibility of any artifacts caused by our ICC method, we verified our results by Western blotting and observed the same rescue effect: Benidipine promoted Miro1 degradation after mitochondrial depolarization in PD fibroblasts (Figure S6C; Figure Legend). We confirmed that Benidipine did not affect *Miro1* messenger RNA (mRNA) expression detected by reverse transcription quantitative real-time PCR (RT-qPCR) under basal and depolarized conditions in PD cells (Figure S6D). Neither did Benidipine alter the basal ATP levels (Figure S6E), nor the mitochondrial membrane potential measured by TMRM staining (Figure S6F).

We next tested Benidipine in the human neuron model of PD described in Figure 3. We confirmed that Benidipine reduced local Ca^2+^ elevation at the OMM detected by GCaMP-Miro1-Anchor (Figure S3F) and promoted Miro1 degradation following Antimycin A treatment in PD neurons (Figure S6G; similar to DFP). This result corroborates that delayed Miro1 removal upon depolarization reflects OMM Ca^2+^ elevation. Notably, treating these PD neurons with Benidipine at 10 μM for 30 hours mitigated stressor-induced neuron death: the loss of TH immunostaining and increase of TUNEL staining after Antimycin A treatment ^33,35^ were significantly rescued (Figure 7A-B).

**Figure 7.**
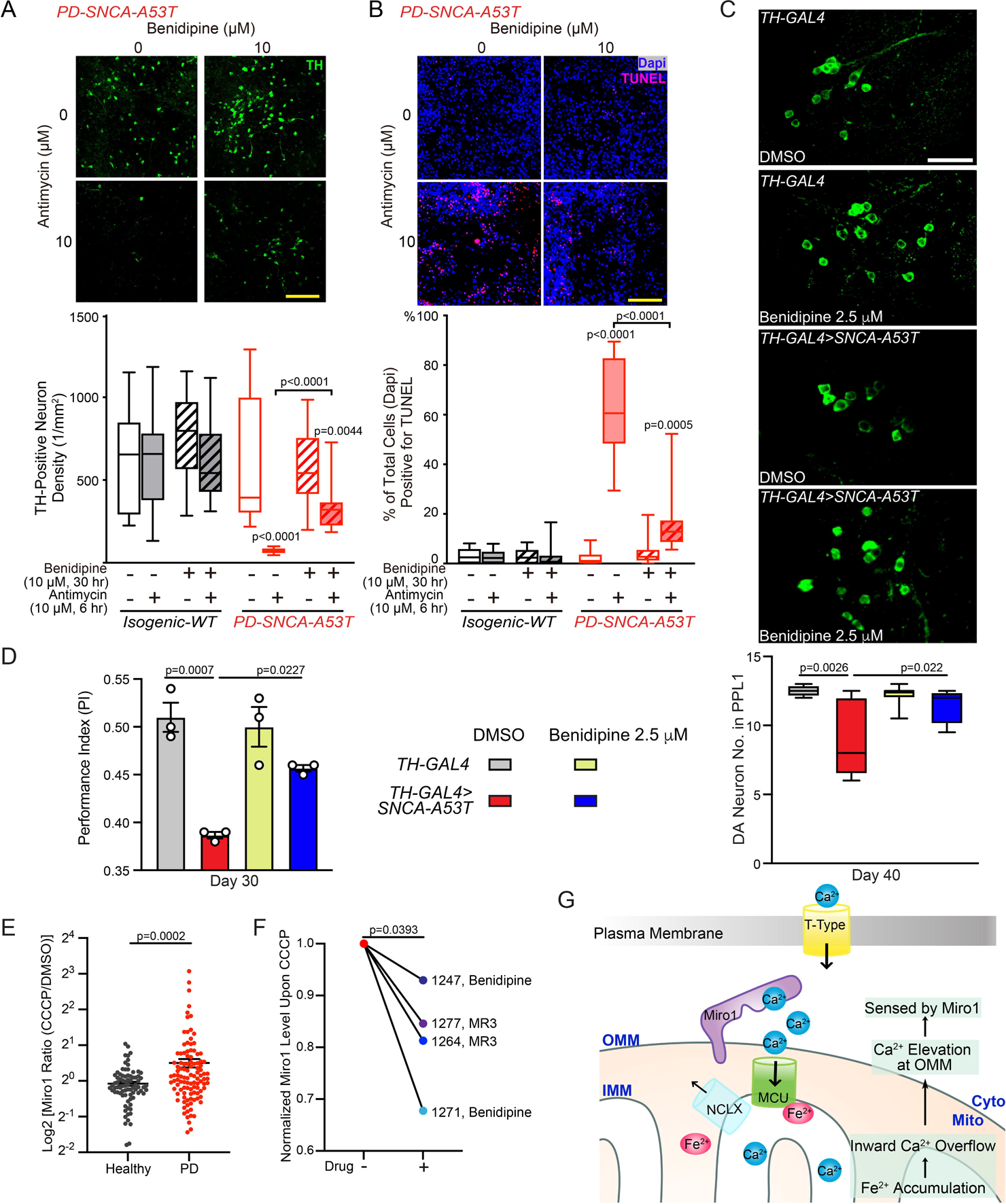
Benidipine Rescues PD Relevant Phenotypes. (A-B) iPSC-derived neurons treated as indicated were immunostained with anti-TH (A) or TUNEL and Dapi (B). Scale bars: 100 μm. n=20 images from 3 independent coverslips. P values are compared with the far-left bar, except indicated otherwise. One-Way Anova Post Hoc Tukey Test. (C) DA neurons in the PPL1 cluster. Scale bar: 20 μm. n=4, 6, 7, 4 (from left to right). (D) n=59, 57, 54, 57 flies (from left to right), 3 independent experiments. (C-D) Drug treatment was started from adulthood (day 1). Two-Way Anova Post Hoc Tukey Test. (E) Miro1 protein levels were measured using ELISA in PBMCs treated with DMSO or 40 μM CCCP for 6 hours. Dot plot with Mean±S.E.M. n=80 controls and 107 PD. Two-tailed Welch’s T Test. (F) PBMCs from 4 PD patients were treated with 40 μM CCCP for 6 hours, or pretreated with 10 μM Benidipine or MR3 for 18 hours and then with 40 μM CCCP for another 6 hours, and Miro1 protein was detected using ELISA. Two-tailed paired T Test. (G) Schematic representation of this study. See also Figure S6-7.

In vivo, we fed Benidipine to the fly model of PD without adding any mitochondrial stressors. Importantly, feeding these flies with 2.5 μM Benidipine from adulthood prevented DA neuron loss in aged flies (Figure 7C) and improved their locomotor ability (Figure 7D). Taken together, our results show that either lowering local OMM Ca^2+^ levels by Benidipine or blocking Miro1’s Ca^2+^ binding eliminates the Miro1 defect of slower degradation upon depolarization and additional phenotypes in PD models (Figure 5-7, S6).

### A Miro1 blood test reflects PD status and responds to drug treatment

A challenge in current PD research is the lack of reliable and convenient molecular markers for patient stratification. The subset of PD patients with Ca^2+^ elevation at the OMM should be sensed by Miro1, and once Miro1 bound to Ca^2+^ it could be quantitatively measured by a proxy–the slower Miro1 response to a mitochondrial uncoupler (Figure 5-7, S6). We next explored the potential of this Miro1 proxy as a PD marker in peripheral blood mononuclear cells (PBMCs). We cultured PBMCs from a healthy donor from the Stanford Blood Center (SBC, Table S4) and depolarized the mitochondrial membrane potential using two different methods: Antimycin A plus Oligomycin ^45^, or CCCP. We found that both depolarizing approaches caused the degradation of Miro1 and additional mitochondrial markers in a time-dependent manner, detected by Western blotting (Figure S7A-B), consistent with other cell types ^32,34,46^. These results show that Miro1 can respond to mitochondrial depolarization in healthy PBMCs, allowing us to further utilize these cells to develop a Miro1 assay for PD.

To enable high-content screening, we applied an ELISA of Miro1 (Figure S7C-D) to PBMCs from the same donor with a 6-hour CCCP treatment. We saw a similar Miro1 response to CCCP using ELISA (SBC, Table S4) as using Western blotting. The reproducibility of the result using ELISA led us to choose this method and 6-hour CCCP treatment to screen a total of 80 healthy controls and 107 PD patients (Table S4). Miro1 Ratio (Miro1 protein value with CCCP divided by that with DMSO from the same person) was significantly higher in PD patients compared to healthy controls (Figure 7E, Table S4), suggesting that PBMCs from more PD patients may have Ca^2+^ elevation at the OMM and consequently fail to rapidly remove Miro1 from damaged mitochondria. Hence, Miro1 Ratio by our method may be used as an indicator of OMM Ca^2+^ changes.

To determine whether our method could be used to classify an individual into a PD or healthy group, we employed machine learning approaches using our dataset. We trained a logistic regression model to assess the impact of Miro1 Ratio on PD diagnosis, solely on its own or combined with additional demographic and clinical parameters (Method). Unified Parkinson’s Disease Rating Scale (UPDRS) is a tool to measure motor and non-motor symptoms of PD which may reflect disease severity and progression. Using UPDRS, our model yielded an accuracy (an individual was correctly classified as with PD or healthy) of 81.2% (p<0.000001; area under the Receiver Operator Curve (ROC)– AUC=0.822), and using Miro1 Ratio, the accuracy was 67.6% (p=0.03; AUC=0.677). Notably, if both Miro1 Ratio and UPDRS were considered, our model generated an improved accuracy of 87.8% (p=0.02; AUC=0.878; p= 5.736e-09 compared to UPDRS alone, paired T test on bootstrapped samples), without the interference of age or sex (Method, Figure S7E-F). Therefore, our results suggest that the molecular (Miro1 Ratio–OMM Ca^2+^ elevation) and symptomatic (UPDRS) evaluations may reveal independent information, and that combining both tests could more accurately categorize individuals with PD and measure their responses to experimental therapies.

To probe the potential utilization of this Miro1 assay in future clinical trials for stratifying patients with OMM Ca^2+^ dysregulation and monitoring drug efficacy, we treated PBMCs from 4 PD patients (Table S4) with either of the two compounds known to reduce Miro1, Benidipine (Figure 6-7, S4-6) and Miro1 Reducer 3 (MR3) ^34,35^. Miro1 protein levels upon CCCP treatment were lowered by each compound in all 4 patients (Figure 7F), showing that the Miro1 marker in PBMCs can respond to drug treatment. Collectively, our results suggest that Miro1 protein in blood cells may be used to aid in diagnosis and drug development.

### Rare variants in T-type Ca^2+^ channels are associated with PD status

After dissecting the functional impairment of this Fe^2+^-Ca^2+^-Miro1 axis in PD, we explored its genetic contribution to PD. Earlier, we showed that chelating iron, blocking MCU’s binding to Fe^2+^, blocking Miro1’s binding to Ca^2+^, or preventing Ca^2+^ entry into the cell all alleviated Parkinson’s related phenotypes (Figures 4-7). We evaluated the genes encoding the protein targets of these approaches, which are spatially distinct and localized to three subcellular locations: (1) IMM Ca^2+^ channels and transporters (targeted by Fe^2+^), (2) the Ca^2+^-binding protein Miro on the OMM, and (3) plasma membrane Ca^2+^ channels (targeted by Benidipine and Tranilast) (Figure 7G, Table S5). By analyzing common variants within or near any of the investigated genes in GWAS reported in ^47^, we did not observe significant association with PD clinical status. We next employed the whole-genome sequencing (WGS) data from the Accelerating Medicines Partnership–Parkinson’s Disease (AMP–PD) (1,168 control; 2,241 PD), and assessed rare non-synonymous and damaging variants using burden based and SKATO methods. We discovered an enrichment of rare variants in selective T-type Ca^2+^-channel subtypes (*Cav3.2*, *3.3*) (cell surface), *Miro2* (OMM), or *NCLX* (IMM) associated with PD status with a nominal P value (Table S5). Notably, a SKATO Test on all variants of T-type or L-type Ca^2+^-channel subtypes showed a significant association with PD status of T-type channels, which survived multiple comparison correction, but not of L-type channels (Table S5). Together, our analysis unravels potential genetic predisposition of T-type Ca^2+^ channels to PD.

### Inhibiting T-type Ca^2+^ channels promotes Miro1 degradation

Our discovery of the selective accumulation of rare variants in T-type Ca^2+^ channels associated with PD diagnosis (Table S5) prompted us to functionally validate this finding. We employed a mini drug screen using the same screening ICC assay described earlier (Figure S4-5) by which we discovered the non-selective pan-Ca^2+^-channel blocker, Benidipine. This time, we examined two L-type and three T-type Ca^2+^-channel blockers. Intriguingly, we again found a striking selection of T-type versus L-type channels, in the connection with Miro1 in PD fibroblasts (Figure S7G): only T-type blockers promoted Miro1 degradation following depolarization, just like Benidipine. Importantly, two of these blockers, MK-8998 ^48, 49^ (FDA approved) and ML-218 ^50^, were already shown in vivo safety and efficacy for treating PD-associated symptoms in humans or preclinical rodent models. By contrast, L-type blockers, including Isradipine that recently failed a phase III trial for PD ^51^, did not affect the Miro1 phenotype (Figure S7G). To genetically inhibit T-type Ca^2+^ channels, we knocked down by siRNA the T-type subtype *Cav3.2* which showed strongest association with PD diagnosis in the human genetic analysis (Table S5). Consistently, genetically reducing *Cav3.2* promoted Miro1 degradation upon depolarization in PD neurons (Figure S7H). These studies implicate a benefit for clinical trials by coupling T-type channel inhibitor drugs and the Miro1 marker.

## Discussion

In this work, we have established a mitochondrial inside-out pathway of Fe^2+^-Ca^2+^-Miro1 dysregulation in our PD models (Figure 7G). Chelating iron, reducing Ca^2+^ entry into the cell, or blocking Miro1’s binding to Ca^2+^ is each neuroprotective (Figure 4, 5, 7). Hence, this ionic axis may be important for PD pathogenesis and can be leveraged for better detecting and treating the disease.

The Miro1 marker could help address unmet needs in PD clinical care. There is a high rate of misdiagnosis of PD because there is no definitive molecular marker to confirm the disease ^52^. The lack of disease-modifying therapies is partially due to no reliable pharmacodynamic biomarkers in clinical trials ^34,53^. Our Miro1 blood test has a potential to serve both as a patient stratification tool and a pharmacodynamic marker for drug discovery. The exciting link between T-type channel blockers and Miro1 in PD suggests this Miro1 blood test may be used in clinical trials for drugs already at the advanced stage, such as MK-8998 ^48,49^. We have also provided evidence that patients’ PBMCs respond to drugs that reduce Miro1. This result opens a door to examining personalized drug efficacy and dosing by testing a patient’s own cells before administrating the drug to the patient, thus providing multiple layers of stratification to improve success rates of clinical trials ^54^. Further studies with independent cohorts are needed to confirm the promise of this test for PD.

Alpha-synuclein aggregation is a pathological hallmark of PD. Emerging evidence has shown that alpha-synuclein has profound functions at the mitochondria or to impact mitochondria, including mitochondrial oxidative phosphorylation, dynamics, mitophagy, Ca^2+^, and iron homeostasis ^7,33,55–60^. Iron accumulation, calcium mishandling, and mitophagy impairment have been individually observed in PD neurons ^2–10,32–34^. However, the relation between these cellular phenotypes in PD has remained obscure. Now we have provided a mechanistic axis connecting them. Fe^2+^ elicits mitochondrial Ca^2+^ overload through acting on IMM Ca^2+^ channels and transporters. The relatively low affinity of MCU to Fe^2+^ in vitro (Figure 2B) is consistent with its low affinity to other divalent metals such as Mg^2+^ ^29^. This data suggests that MCU may not be predominantly occupied by these divalent ions under physiological conditions ^29^. By contrast, under pathological conditions with elevated Fe^2+^, MCU binds to more Fe^2+^ and cellular detriment ensues. This finding agrees with prior evidence showing that inhibiting MCU prevents Fe^2+^-induced cell death ^36,61^ and Parkinsonian neurodegeneration ^1,8^. Further investigations are needed to dissect how Fe^2+^ regulates MCUb and NCLX levels. Our results (Figure 2G-H) have suggested that Fe^2+^-triggered Ca^2+^ efflux delay and the reduction of NCLX and MCUb depend on mitochondrial Ca^2+^ overload. Thus, it is possible that NCLX and MCUb are targeted by Ca^2+^-activated mitochondrial proteases. Our data also suggests Miro1 may link Ca^2+^ and mitophagy phenotypes in PD. In the same PD neuron models used in this paper, we have previously characterized a delayed mitophagy phenotype downstream of Miro1 retention on damaged mitochondria ^32–34^. Our paper now shows this Miro1 phenotype is dependent on OMM Ca^2+^ elevation because of iron accumulation. In addition to mitophagy ^32,41,62^, Miro may have additional functions in regulating mitochondrial quality control, including the biogenesis of mitochondrial derived vesicles ^63^ and transcellular mitochondrial transfer ^64–68^. When Miro senses spatially restricted Ca^2+^ elevation at the OMM in PD neurons, it may affect multiple Miro-mediated biological processes. More studies are warranted to unravel the precise roles of Miro in PD pathogenesis and how Ca^2+^ regulates these roles.

### Limitation of the study

Flies do not contain an alpha-synuclein homolog, and our results may not account for the physiological roles of human alpha-synuclein. The sample size for the postmortem brain study is relatively small and a larger cohort of PD and PD-related disorders is needed to confirm the prevalence and specificity of the phenotype. Re-evaluation of genetic variants in this pathway is warranted in additional larger cohorts to verify their link to PD. Although MCU oligomerization could be enhanced with a micromolar range of Fe^2+^ added in the media (Figure 1F), the pathophysiological level of free Fe^2+^ is difficult to determine in vivo.

## Acknowledgements

We thank M.F. Tsai for *MCU* KO cells, L. Pallanck for flies, A.Y. Ting for constructs, and P. Katiyar, Z.T. Cook, D.M. Conradson, and J. Panji for technical support, Stanford SIGMA Facility for ICP-MS (NSF ECCS-2026822), Stanford HTBC (RRID: SCR_017794) and D. E. Solow-Cordero for drug screens, and SBC, Stanford ADRC, UCLA Department of Pathology, R.N. Alcalay, Columbia University, and Banner Sun Health Research Institute for providing human samples. This work was supported by NINDS (RO1NS089583, RO1NS128040; X.W.), NIGMS (RO1GM143258; X.W.), NIA (RO1AG060747; M.D.G.), Stanford SPARK and Spectrum SPADA (X.W.), Sanofi iAward (X.W.), Warren Alpert Foundation (X.W.), MJFF (021146; X.W.), Harrington Discovery Institute (X.W.), Belgian American Education Foundation (R.V.), Glenn Foundation Postdoctoral Fellowship (L.L.), and European Union’s Horizon 2020 research and innovation program under Marie Skłodowska-Curie (890650; Y.LG.). Banner Sun Brain and Body Donation Program was supported by NINDS (U24 NS072026), NIA (P30 AG19610), Arizona Department of Health Services (211002), Arizona Biomedical Research Commission (4001, 0011, 05-901 and 1001), and MJFF.

## Author contributions

V.B., A.S.D., R.V., C.-H.H., S.C., L.L., and C.S.K. performed human cell experiments. R.V. did drug screens. L.L. conducted fly work. C.M.M. purified MCU and did FSEC with assistance from L.F. C.S.K. did molecular cloning. Y.LG. and M.D.G. analyzed human genetic data. P.N. analyzed PBMC data. X.W. conceived and supervised the project. V.B., R.V., and X.W. wrote the paper with assistance from all authors.

## Inclusion and diversity

We support inclusive, diverse, and equitable conduct of research.

## STAR METHODS

### Resource availability

#### Lead contact

Further information and requests for resources and reagents should be directed to the lead contact Xinnan Wang.

#### Materials availability

All reagents generated in this study are available from the lead contact.

#### Data and code availability

- All original imaging and Western blotting data are available from the lead contact upon request.
- Original code and additional method for machine learning has been deposited at Mendeley and is publicly available as of the date of publication. DOI is listed in Method and Key Resource Table.
- Any additional information required to reanalyze the data reported in this paper is available from the lead contact upon request.

### Experimental model and study participant details

#### Human cells and tissues

No human subjects were used in this study. The iPSC work was approved by Stanford Stem Cell Oversight Committee. iPSCs or fibroblasts were purchased under a material transfer agreement (MTA) from NINDS Human Cell and Data Repository or Jackson Laboratory. PBMCs were obtained under an MTA from Columbia University and Michael J. Fox Foundation (MJFF), and from SBC and Stanford Alzheimer’s Disease Research Center (ADRC). Postmortem brain tissues were obtained from Stanford ADRC, Department of Pathology at University of California Los Angeles (UCLA), and Banner Sun Health Research Institute. Columbia University, Stanford University, UCLA, and Banner Sun Health Research Institute Institutional Review Board approved study protocols, which ensured consent from donors and explained the conditions for donating materials for research. Demographic and clinical details of each subject are in Supplementary Tables and Key Resource Table. HEK293T cells were obtained from ATCC. Details of cell culture conditions and authentication are in Method Details.

#### Fly Model

The following fly stocks were used: *Actin-GAL4* (Bloomington Drosophila Stock Center–BDSC), *TH-GAL4* (BDSC), *UAS-SNCA^A53T^*^69^, and *UAS-T7-DMiro-WT* ^70^. *UAS-T7-DMiro-KK* were generated by injecting pUAST-T7-DMiro-KK into flies by Bestgene Inc (Chino Hills, CA). All fly stocks and experiments were kept at 25°C with a 12:12 hrs light:dark cycle and constant humidity (65%) on standard sugar-yeast-agar (SYA) medium (15 g/l agar, 50 g/l sugar, 100 g/l autolyzed yeast, 6 g/l nipagin, and 3 ml/l propionic acid) ^71^. Flies were raised at standard density in 200 ml bottles unless otherwise stated. All fly lines were backcrossed 6 generations into a *w*^1118^ background to ensure a homogeneous genetic background. The ages of flies for each experiment were stated in figures. Experiments were carried out on mated females ^72^.

### Method details

#### Constructs

pcDNA3.1-MCU-Flag was purchased from Genescript (MCU_OHu20123D_pcDNA3.1+/C-(K)-DYK). pcDNA3.1-MCU-3A-Flag was generated by Synbio Technologies by making 3 point mutations (74D, 148D, 159H to A) of pcDNA3.1-MCU-Flag. pRK5-GFP-hMiro1-WT or KK was custom-made by Synbio Technologies. pDsRed2-Mito was described in ^32^. pUAST-T7-DMiro-KK was generated by introducing E234K, E354K to pUAST-T7-DMiro ^70^. Myc-hMiro1-WT and was described in ^43^. pLX304-miRFP670nano_P2A_GCaMP6f-Miro1-Anchor or pLX304-miRFP670nano_P2A_ER-GCaMP6f (cytosol) was made by cutting pLX304-miRFP670nano_P2A_EGFP (a gift from Alice Y. Ting) with BsiW1 and Nhe1, amplifying each gene fragment by PCR, and then ligating these with the Gibson assembly method (HiFi master mix, NEB). pcDNA3.1-ER-GCaMP6f (cytosol) was a gift from Chris Richards (Addgene #182548). miRFP670nano was a gift from Vladislav Verkhusha (Addgene #127443). All vectors were confirmed by sequencing.

#### Cell Culture

Human dermal primary fibroblasts isolated from a PD patient (ND39528) and a healthy control (ND36091) were cultured in DMEM (11995-065, Thermo Fisher) supplemented with 10% fetal bovine serum (900-108, heat-inactivated, Gemini Bio-Products), 1×Anti Anti (15240096, Thermo Fisher), and 1×GlutaMax (35050061, Thermo Fisher) maintained in a 37°C, 5% CO_2_ incubator with humidified atmosphere. HEK293T cells were cultured in DMEM (11995-065, Thermo Fisher) supplemented with 10% fetal bovine serum (900-108, heat-inactivated, Gemini Bio-Products), 1% Pen-Strep (15140-122, Gibco), and maintained in a 37°C, 5% CO_2_ incubator with humidified atmosphere. Different concentrations of Fe^2+^ (Ammonium Iron(II) Sulfate Hexahydrate, 203505, Sigma-Aldrich) were added to the media for 20-24 hours. The MTT assay (ab211091, Abcam) was used according to the manual. Cells were treated with 10 μM Benidipine (B6813, Sigma-Aldrich), 40 μM CCCP (C2759, Sigma-Aldrich), 100 μM VitC (A5960, Sigma-Aldrich), or 100 μM DFP (HY-B0568, MedChemExpress).

#### iPSC Culture and Transfection

All iPSC lines in this study were fully characterized and authenticated by our previous studies ^32,33^, NINDS Human Cell and Data Repository (https://stemcells.nindsgenetics.org/), Applied StemCell, or Jackson Laboratory (https://www.jax.org/jax-mice-and-services/ipsc/cells-collection). iPSCs were derived to midbrain DA neurons as previously described with minor modifications ^32,73–76^. Briefly, neurons were generated using an adaptation of the dual-smad inhibition method. Neural induction was started when iPSCs reached 50-85% confluency, with the use of dual smad inhibitors, dorsormorphin (P5499, Sigma-Aldrich) and TGFβ inhibitor SB431542 (1614, Tocris), and the addition of GSK3β inhibitor CHIR99021 (04-0004, Stemgent) and smoothened agonist SAG (566661, CalBioChem). To get single cells, 12 days after neural induction, half of the medium was replaced with N2 medium with 20 ng ml^−1^ BDNF (450-02, Peprotech), 200 μM ascorbic acid (A5960, Sigma-Aldrich), 500 nM SAG, and 100 ng ml^−1^ FGF8a (4745-F8-050, R&D Systems). On day 16-17, neurons were split and transferred onto Matrigel (354277, Corning)-coated or poly-ornithine and laminin-coated glass coverslips in a 24-well plate. On day 19-20, medium was switched to N2 medium supplemented with 20 ng ml^−1^ BDNF, 200 μM ascorbic acid, 20 ng ml^−1^ GDNF (450-10, Peprotech), 1 ng ml^−1^ TGFβ3 (AF-100-36E, Peprotech), and 500 μM dibutyryl-cAMP (D0627, Sigma-Aldrich) for maturation of DA neurons. Neurons were used at day 21-26 after neuronal induction in most of the experiments, when about 80-90% of total cells expressed the neuronal marker TUJ-1, and 14-17% of total cells expressed TH and markers consistent with ventral midbrain neuronal subtypes ^33,77^. For transfection, on day 19-20 after neural induction, culture medium was replaced with Opti-MEM (Gibco) prior to transfection. For 24-well plates, 0.5-1 μg DNA or 1 μl Lipofectamine 2000 (11668-030, Invitrogen) was diluted in Opti-MEM at room temperature (22°C) to a final volume of 50 μl in two separate tubes, and then contents of the two tubes were gently mixed, incubated for 20 minutes at room temperature, and subsequently added onto neurons (75,000-120,000 per well). For 6-well plates, 1.5 μg DNA or 6 μl Lipofectamine 2000 (11668-030, Invitrogen) was diluted in Opti-MEM to a final volume of 250 μl in each tube, and mixed, incubated, and applied to neurons (700,000-1,500,000 per well). After 6 hours, Opti-MEM containing DNA-Lipofectamine complexes was replaced with regular N2 medium with supplements. For siRNA, NT siRNA (D-001910-10, Horizon Discovery), *Cav3.2* siRNA (EQ-006128-00-0002, Horizon Discovery), and delivery media were used according to the manual. After transfection for 2-3 days, neurons were used. Neurons from 6-well plates were lysed in 400 μl lysis buffer (300 mM NaCl, 50 mM Tris pH 7.5, 1% Triton X-100, 0.2 mM PMSF, Protease Inhibitor Cocktail, 5 mM EDTA) and used in the Miro1 ELISA kit (EKL54911, Biomatik) following the manual, or in 300 μl Extraction Buffer supplemented with 0.2 mM PMSF and Protease Inhibitor Cocktail and used in the GFP ELISA kit (ab229403, Abcam) following the manual. Neurons were treated with 50-100 μM DFP (HY-B0568, MedChemExpress), 10 μM Benidipine (B6813, Sigma-Aldrich), 500 nM RU360 (557440, Sigma-Aldrich), 10 μM Antimycin A (A8674, Sigma-Aldrich), or 100 μM Antimycin A during live imaging.

#### Primary Drug Screens

Overall scheme: Fibroblasts were plated and cultured for 24 hours. Compound libraries were then applied for 10 hours. Next, 20 µM FCCP was added for another 14 hours. FCCP is a mitochondrial uncoupler that depolarizes the mitochondrial membrane potential ^78^. Application of mitochondrial uncouplers had been successfully used in high-throughput screens searching for genes and chemicals in the mitophagy pathway ^79–81^. After fixation, cells were stained with anti-Miro1 and 4’,6-diamidino-2-phenylindole, dihydrochloride (Dapi), and imaged under the confocal microscope with the identical setting (Figure 6A). Anti-Miro1 has been validated in ICC for specifically recognizing endogenous Miro1 using human fibroblast-derived neurons with *Miro1* RNAi ^32^. The Miro1 intensity was normalized to Dapi from those images. Using this analysis, we observed significant Miro1 reduction following FCCP treatment in fibroblasts from a healthy control subject but not in a sporadic PD fibroblast line (Figure S6A), consistent with our previous studies using the same cell lines by Western blotting or ELISA ^34^. We then employed this method to screen 3 chemical libraries (NIH, FDA, and Sigma) using the PD cell line. The NIH Clinical collection library contains 377 unique compounds. We screened those drugs at a defined concentration of 10 µM with 4 biological repeats (independent wells) for each drug. We compared calculated Miro1 levels in PD cells treated with both compound and FCCP to those treated with FCCP but without any compound. We found that 11 out of 377 unique compounds reduced Miro1 protein levels following FCCP treatment on or below 3SD (standard deviation) of control Miro1 values (those from the same PD cell line without compound treatment but with FCCP), in all biological repeats (Figure S4A, Table S2A) ^82^. We next screened the Biomol FDA library with two five-fold doses (1.25, 2.5, 5, 10, 20 µM). From 175 unique compounds, we identified 3 hits that reduced Miro1 protein levels following FCCP treatment in a dose-dependent manner (Figure S4B, Table S2A). Lastly, we screened the Sigma LOPAC library which contains 1269 unique compounds at a defined concentration of 20 µM. We found that 21 compounds reduced Miro1 protein levels on or below 3SD of those control Miro1 values from the same PD cell line without compound treatment but with FCCP (Figure S4C, Table S2A). We ranked all hits in the order of the degree of Miro1 reduction (Table S2A). It is important to note that from the same screens we also identified compounds that enhanced Miro1 protein levels in PD cells treated with FCCP (Table S2B). Although our purpose was to search for Miro1 reducers in PD models, the same assay could be used to look for Miro1 enhancers in other disease models.

Method details: Fibroblasts were plated onto clear-bottomed/black-walled 384-well plates (EK-30091, E&K Scientific) at 2,000 cells/well with a Matrix Wellmate dispenser (Thermo Fisher), and plates were incubated at 37°C, 5% CO_2_ for 24 hours. Next, chemical library compounds at defined concentrations were added using fully automated liquid handling system (Caliper Life Sciences Staccato system) with a Twister II robot and a Sciclone ALH3000 (Caliper Life Sciences) integrated with a V&P Scientific pin tool, and plates were incubated at 37°C, 5% CO_2_ for 10 hours. Then, 20 µM of FCCP (C2920, Sigma-Aldrich) was added to wells and plates were incubated at 37°C, 5% CO_2_ for another 14 hours. Cells were fixed with ice-cold 90% methanol (482332, Thermo Fisher) for 20 minutes at -20°C, then incubated with blocking buffer (10% normal goat serum–50062Z, Thermo Fisher; 0.5% BSA–BP1600, Thermo Fisher; 0.2% Triton X-100–T8787, Thermo Fisher) at room temperature for 15 minutes, and incubated with anti-Miro1 (HPA010687, Sigma-Aldrich) at 1:100 in blocking buffer overnight at 4°C. Samples were washed with 1×PBS (10010-049, Thermo Fisher) using a Plate Washer multivalve (ELx405UV, Bio-Tek), incubated with goat anti-Rabbit IgG (H+L) Cross-Adsorbed, Alexa Fluor 488 (A11008, Thermo Fisher) at 1:500 in blocking buffer at 25°C for 2 hours, then washed again with 1×PBS, and finally 1.0 µg/ml Dapi (D1306, Thermo Fisher) in 1×PBS was added and plates were sealed using PlateLoc (01867.001, Velocity11). All liquids were dispensed using the Multidrop 384 (5840200, Titertek) unless otherwise stated. Fluorescent signals in plates were automatically imaged with ImageXpress Micro (Molecular Devices, IXMicro) and data were analyzed with MetaXpress Analysis (Molecular Devices). One image was taken from one well. Our negative controls without primary anti-Miro1 generated no Miro1 signals. Only cells positive for both Miro1 and Dapi staining were chosen for analysis (>99%; total 400-600 cells per image). The Miro1 fluorescent intensity was normalized to that of Dapi from the same cell and averaged across all cells from the image from one well. Then, the median of the mean Miro1 intensities of all wells from the same plate was calculated and subtracted from the mean Miro1 intensity of each well.

#### Retesting Hits from the Primary Screens

##### Overall scheme

To validate the results of the high-content assays, we retested 34 out of the 35 positive Miro1 reducers identified at the Stanford HTBC in our own laboratory using fresh compounds and our confocal microscope. We didn’t test 1 hit (CV-3988) because it was unavailable for purchase at the time of the experiments. We applied each of the 34 compounds at the highest screening concentration to the same sporadic PD line used at the HTBC with at least 4 biological replicates. We also evaluated Miro1 protein levels without FCCP but with compound treatment, which unveiled the drug effect on Miro1 under the basal condition in PD fibroblasts. We imaged all samples with the identical imaging setting, and our negative controls without primary anti-Miro1 yielded almost no signals (background; Table S2C). Four compounds, A-77636 hydrochloride, Ebselen, GBR-12909 dihydrochloride and Fenbufen, appeared toxic (Table S2C). We found that 15 out of the initial 34 hit compounds consistently reduced Miro1 protein following FCCP treatment below 2SD of control Miro1 values (those from the same PD cell line without treatment) (Figure S5, Table S2C).

Method details: Fibroblasts were plated on coverslips (22-293232, Fisher scientific) into 24-well plates (10861-558, VWR) at 20,000 cells/well and plates were incubated at 37°C, 5% CO_2_ for 24 hours. Next, fresh chemical compounds dissolved in DMSO were added at defined concentrations and plates were incubated at 37°C, 5% CO_2_ for 10 hours. Then, FCCP at 20 µM was added to wells and plates were incubated at 37°C, 5% CO_2_ for another 14 hours. Cells were fixed with ice-cold 90% methanol for 20 minutes at -20°C, incubated with blocking buffer (10% normal goat serum, 0.5% BSA, 0.2% Triton X-100) at 25°C for 15 minutes, and immunostained with anti-Miro1 (HPA010687, Sigma-Aldrich) at 1:100 in blocking buffer overnight at 4°C. Samples were washed three times with 1×PBS, incubated with goat anti-Rabbit IgG (H+L) Cross-Adsorbed, Alexa Fluor 488 (A11008, Thermo Fisher) at 1:500 in blocking buffer at 25°C for 2 hours, washed again three times with 1×PBS, and finally samples were mounted with ProLong™ Glass Antifade Mountant with NucBlue™ (Dapi) Stain (P36983, hard-setting, Thermo Fisher) on glass slides and allowed to cure overnight. Samples were imaged at 25°C with a 20×/N.A.0.60 oil Plan-Apochromat objective on a Leica SPE laser scanning confocal microscope (JH Technologies), with identical imaging parameters among different genotypes. Three images were taken per coverslip. Total 4 biological repeats (coverslips) for each drug. Images were processed with ImageJ (Ver. 1.48, NIH), and the average Miro1 intensity in the cytoplasmic area was measured using the Intensity Ratio Nuclei Cytoplasm Tool (http://dev.mri.cnrs.fr/projects/imagej-macros/wiki/Intensity_Ratio_Nuclei_Cytoplasm_Tool). All compound information for retesting is in Table S2C or in Figure S7.

#### Pathway Analysis

Putative mechanistic pathways were generated using graph search over a network of over 2 million relationships between chemicals, diseases, and genes extracted from the abstracts of journal publications deposited in PubMed ^83^. Each chemical hit and Miro1 were used as the source and target inputs for a weighted shortest path algorithm where paths were only allowed to traverse chemical-gene and gene-gene relationships. Graph search over the biomedical graph was performed using Neo4J. A subnetwork associated with each shortest path was generated from its supporting documents and analyzed using a knowledge graph browser (docs2graph) to curate the set of supporting documents.

#### Immunocytochemistry and Confocal Microscopy

Adult fly brains were dissected in PBST (0.3% Triton X-100 in PBS) and incubated with fixative solution (4% formaldehyde in PBST) for 15 minutes at room temperature. Fixed samples were incubated in blocking solution (10% BSA in PBST) for an hour, and then immunostained with rabbit anti-TH (AB-152, EMD Millipore) at 1:200 in blocking solution for 36 hours at 4°C on a rotator. After wash, donkey anti-rabbit Alexa Fluor 568 (ab175470, Abcam) at 1:1000 was applied in blocking solution for 1 hour at room temperature in dark. The DA neuron number was counted throughout the Z stack images of each brain. Representative images were summed stack images. Human neurons were fixed in 4% paraformaldehyde (Electron Microscopy Sciences, Hatfield, PA) for 15 minutes at room temperature, permeabilized with 0.25% Triton X-100 in PBS for 20 minutes, then washed twice in PBS (5 minutes each) and rinsed in deionized water. Alexa Fluor 594 picolyl azide based TUNEL assay was performed according to the manufacturer’s instruction (C10618, Invitrogen). Coverslips were incubated with TdT reaction buffer at 37°C for 10 minutes, followed by TdT reaction mixture at 37°C for 60 minutes in a humidified chamber. Coverslips were then rinsed with deionized water, washed with 3% BSA and 0.1% Triton X-100 in PBS for 5 minutes, followed by incubation with Click-iT plus TUNEL reaction cocktail at 37°C for 30 minutes, and then washed with 3% BSA and 0.1% Triton X-100 in PBS for 5 minutes. For nuclear counterstain, 4’, 6-Diamidino-2-phenylindole (Dapi; D9542, Sigma-Aldrich) at 0.5 µg/ml was applied at room temperature for 10 minutes in the dark. The coverslips were washed with PBS twice, and then mounted on slides with Fluoromount-G mounting medium (SouthernBiotech). For some experiments, Dapi was stained with ProLong™ Glass Antifade Mountant with NucBlue™ Stain (P36983, hard-setting, Thermo Fisher). Or iPSC-derived neurons were fixed in 4% paraformaldehyde for 15 minutes, washed twice in PBS (5 minutes each), and then blocked in PBS with 5% normal goat serum and 0.3% Triton X-100 for 60 minutes. Neurons were then immunostained with rabbit anti-TH (AB-152, EMD Millipore) at 1:500, rabbit anti-Flag (F7425, Sigma-Aldrich) at 1:500, or mouse anti-Myc at 1:1,000 (sc-40, Santa Cruz) in antibody buffer (PBS with 1% BSA and 0.3% Triton X-100) at 4°C overnight, followed by Alexa Fluor 488 fluorochrome conjugated goat anti-rabbit IgG (A11008, Invitrogen) or Alexa Fluor 568 fluorochrome conjugated goat anti-mouse IgG (A11004, Invitrogen) at 1:500 in antibody buffer at room temperature for 1 hour. Coverslips were washed with PBS twice, and then mounted on slides with Fluoromount-G mounting medium. Samples were imaged at room temperature with a 20×/N.A.0.60 or 63×/N.A.1.30 oil Plan-Apochromat objective on a Leica SPE laser scanning confocal microscope, with identical imaging parameters among different genotypes in a blinded fashion. Images were processed with ImageJ (Ver. 1.48, NIH) using only linear adjustments of contrast and color.

#### Live Cell Imaging

RPA at 5 μM (ME043.1, Squarix biotechnology), Mito-FerroGreen at 5 μM (M489, Dojindo Laboratories), MitoTracker Green at 75-100 nM (M7514, Thermo Fisher), or TMRM at 25 nM (T668, Molecular Probes) was applied to cells on coverslips in culture media or Hanks Balanced Salt Solution (HBSS) at 37°C for 30 minutes, then coverslips were washed with HBSS and each was placed in a 35-mm petri-dish containing Hibernate E low-fluorescence medium (HELF, BrainBits/Transnetyx), and cells were imaged immediately on a heated stage at 37°C with a 63/N.A.0.9 water-immersion objective. For calcium imaging, Rhod-2 (R1244, Thermo Fisher) and Calcium Green (C3011MP, Thermo Fisher) at 5 μM were applied to cells on coverslips and incubated at 37°C for 30 minutes and imaged with the same set-up as above. Thrombin (10602400001, Sigma-Aldrich) was applied at 100 mUnits/ml to the dish during imaging. 500 μM Fe^2+^ was applied the day before. Time-lapse movies were obtained continually with a 2 second interval for 8-10 minutes. For Mito-dsRed and GFP-Miro1 imaging, live neurons were imaged as above and time-lapse movies were obtained continually with a 1 second interval before and after 100 μM Antimycin (A8674, Sigma-Aldrich) addition.

#### Fly Behavioral Analysis

DFP (HY-B0568, MedChemExpress) was dissolved in water at 50 mM and added to SYA fly food at a final concentration of 163 μM. Benidipine (B6813, Sigma-Aldrich) was dissolved in DMSO at 10 mM and added to SYA fly food at a final concentration of 2.5 μM. The same volume of solvent was added to control food. Drug treatment was started on newly-eclosed mated females (day 1), except for DFP on locomotion which was started from embryogenesis. Food was refreshed every 2 days. The average Performance Index (PI) (negative geotaxis) was evaluated as previously described ^72,84,85^. Briefly, adult flies were gently tapped to the base of a modified 25 ml climbing tube and their climbing progress was recorded after 45 seconds. Two to four populations of flies were assessed, and for each population, flies were examined 3 times per experiment. The recorded values were used to calculate the average PI.

#### Western Blotting and IP

In general, lysates for SDS-PAGE were mixed with 4×Laemmli sample buffer (161-0747, Bio-Rad) containing 5-10% β-mercaptoethanol, and then boiled for 5Lminutes before being loaded into either a precast 10% polyacrylamide gel (4561034, Bio-Rad) or manually made gel for electrophoresis with Tris-glycine-SDS running buffer (24.8LmM Tris, 192LmM glycine, 0.1% SDS). For Native-PAGE, cells were lysed in lysis buffer (300 mM NaCl, 50 mM Tris pH 7.5, 1% Triton X-100, 0.5% Digitonin, 5 mM EDTA, 0.2 mM PMSF, Protease Inhibitor Cocktail–P8340, Sigma-Aldrich), and centrifuged at 16,200 g 4°C for 1 hour. Lysates were mixed with 2×Native Tris-Glycine sample buffer (LC2673, Thermo Fisher) and loaded directly into a precast 10% polyacrylamide gel (4561034, Bio-Rad) with Tris-glycine Native running buffer (24.8LmM Tris, 192LmM glycine). After electrophoresis, nitrocellulose membranes (1620115, Bio-Rad) were used in semi-dry transfer with Bjerrum Schafer– Nielsen buffer (48LmM Tris, 39LmM glycine, 20% methanol (v/v), pH 9.2). Transferred membranes were first blocked in TBST (TBS with 0.01-0.05% Tween-20) with 5% milk for 1Lhour at room temperature, and then immunoblotted with the following primary antibodies in TBST with 5% milk at 4°C overnight: mouse anti-Miro1 (WH0055288M1, Sigma-Aldrich) at 1:1,000, rabbit anti-GAPDH (5174S, Cell Signaling Technology) at 1:1,000, rabbit anti-MCU (HPA016480, Sigma-Aldrich) at 1:500, rabbit anti-Flag (F7425, Sigma-Aldrich) at 1:750-1,000, rabbit anti-NCLX (ab136975, Abcam) at 1:1,000, rabbit anti-MCU (26312-1-AP, Proteintech) at 1:1,000, rabbit anti-Miro2 (14016S, Cell Signaling Technology) at 1:1,000, mouse anti-ATP5β (ab14730, Abcam) at 1:1,000, rabbit anti-MCUb (AP12355b, Abgent) at 1:1,000, rabbit anti-β-Actin (4967S, Cell Signaling Technology) at 1:1,000, rabbit anti-Mitofusin2 (9482S, Cell Signaling Technology) at 1:1,000, mouse anti-MIC60 (ab110329, Abcam) at 1:500, mouse anti-MIC19 (TA803454, Origene) at 1:1,000, rabbit anti-MICU1 (HPA037479, Sigma-Aldrich) at 1:1,000, rabbit anti-MICU2 (ab101465, Abcam) at 1:1,000, rabbit anti-FECH (14466-1-AP, Proteintech) at 1:500, rabbit anti-EMRE (HPA032117, Sigma-Aldrich) at 1:500, rabbit anti-OPA1 (80471S, Cell Signaling Technology) at 1:1,000, mouse anti-T7 (69622, Novagen) at 1:1,000, and rabbit anti-VDAC (4661S, Cell Signaling Technology) at 1:1,000. Secondary HRP-conjugated goat anti-rabbit or mouse IgG antibodies (Jackson ImmunoResearch Laboratory) were used at 1:3,000–10,000 for 1 hour at room temperature. West Dura ECL Reagent (34075, GE Healthcare) or SuperSignal^TM^ West Pico PLUS Chemiluminescent Substrate (Thermo fisher scientific) was added. Membranes were scanned using a Bio-Rad ChemiDoc XRS system.

HEK cells (0.3×10^6^/well in a 6-well plate) were plated, grown overnight, and next day when 30-40% confluent, transfected with 1 μg of DNA, using the calcium phosphate transfection protocol. After 24 hours of transfection, media was changed. After 48 hours of transfection, cells were treated without or with 500 μM Fe^2+^ (Ammonium Iron(II) Sulfate Hexahydrate, 203505, Sigma-Aldrich) for 24 hours followed by lysing with 300 μl of lysis buffer (300 mM NaCl, 50 mM Tris pH 7.5, 1% Triton X-100, 0.2 mM PMSF, Protease Inhibitor Cocktail) supplemented with 5 mM EDTA or 500 μM Fe^2+^. For each condition, 3 wells of a 6-well plate were pooled, followed by centrifugation at 16,200 g for 10 minutes at 4°C. Part of the supernatant (110 μl out of 900 μl total) was reserved as ‘Input’. The remaining 790 μl was incubated with 10 μl of anti-Flag (F7425, Sigma-Aldrich) for 2 hours on a rotator at 4°C, and then combined with 60 μl of washed Protein A–Sepharose beads (17-0780-01, GE Healthcare) for 1 hour on a rotator at 4°C. Beads were then washed five times with lysis buffer. Residual buffer was removed from the last wash and the beads were mixed with 100 μl of 2×Tris-Glycine Native Sample Buffer (LC2673, Thermo Fisher) or Laemmli buffer (then boiled), and loaded into a precast gel (4561034, Bio-Rad). For each gel running, approximately 5% (45 μl out of total 900 μl lysate) of total lysates (Input) and 45% of total immunoprecipitated proteins were loaded.

#### Postmortem Brain Lysing for Western Blotting

A piece of postmortem brain (1 mm^3^) was cut and put in a 1.5 ml tube, and lysed in 500 μl of lysis buffer (300 mM NaCl, 50 mM Tris pH 7.5, 1% Triton X-100, 0.5% Digitonin, Protease Inhibitor Cocktail, 0.2 mM PMSF). Tissue was sonicated and then centrifuged at 16,200 g 4°C for 1 hour. Supernatant was collected.

#### RT-qPCR

Total RNA was extracted from at least one million cells per sample using TRIzol (Gibco) according to the manufacturer’s instructions. Concentrations of total RNA were measured using a NanoDrop. 1 μg of total RNA was then subjected to DNA digestion using DNase I (Ambion), immediately followed by reverse transcription using the iScript Reverse Transcription Supermix (1708841, BIO-RAD). qPCR was performed using the StepOnePlus^TM^ instrument (Thermo Fisher) and SYBR Green Supermix (172-5270, Bio-Rad) following the manufacturer’s instructions. Human *GAPDH* was amplified as an internal standard. Expression levels were analyzed by the StepOne Software (Version 2.2.2). The relative expression level was divided by that of *GAPDH* from the same experiment. Each experiment was repeated 3-4 times in duplicate.

#### ATP Levels

ATP concentrations were determined using the Roche ATP Bioluminescence Assay Kit HS II (no. 11699709001, Sigma). Briefly, cell lysate was homogenized in 150 μl of ice-cold lysis buffer as above using a Kontes pellet pestle. Lysate was then boiled for 5 minutes and centrifuged at 20,000 g 4°C for 1 minute. Cleared lysate was diluted 1:200 in dilution buffer and loaded with 10 μl of luciferase. Luminescence was immediately measured using a FlexStation 3 (Molecular Devices). Total protein amounts were measured using the Bicinchoninic Acid (BCA) Protein Assay Kit (23227, Thermo Scientific). The ATP level in each sample was normalized to the total protein amount.

#### SEC

For control and H_2_O_2_ treatment (10 mM for 10 minutes), HEK cells were lysed in lysis buffer (1% Triton X-100, 50 mM Tris pH7.5, 300 mM NaCl, 5 mM EDTA, 1:1000 Protease inhibitor cocktail, 0.2 mM PMSF). For Fe^2+^ treatment, HEK cells were treated with 5 mM Fe^2+^ (Ammonium Iron(II) Sulfate Hexahydrate, 203505, Sigma-Aldrich) for 24 hours, and lysed in the same buffer above where EDTA was replaced with 5 mM Fe^2+^. Cell lysates were injected onto a Superdex 200 Increase 10/300 GL column (GE Healthcare Life Sciences). To slow Fe^2+^-oxidation and precipitation in equilibration buffer, we preequilibrated the column with low pH buffer and reducing reagent. For the control sample, the column was preequilibrated with 20 mM MES pH 6.0, 300 mM NaCl, 5 mM TCEP, and 0.1% Triton X-100. For the Fe^2+^-treated sample, the column was preequilibrated with 20 mM MES pH 6.0, 300 mM NaCl, 5 mM TCEP, 2 mM Fe^2+^, and 0.1% Triton X-100. For the H_2_O_2_-treated sample, the column was preequilibrated with 20 mM Tris pH 7.4, 300 mM NaCl, and 0.1% Triton X-100.

Fluorescence SEC: For each sample, 5 µg purified human MCU was incubated either on ice or at 35°C in a thermal cycler for 15 minutes. After the incubation, all samples were centrifuged in a tabletop centrifuge at 21,000 g at 4°C for 10 minutes to remove precipitated protein. The supernatant was then injected onto a Superdex 200 Increase 10/300 GL column preequilibrated with 20 mM Tris pH 7.4, 150 mM NaCl, 0.01% lauryl maltose neopentyl glycol (LMNG), and 0.001% cholesteryl hemisuccinate (CHS).

#### Iron Detection Assay

HEK cells from one well of a 6-well plate were lysed in 300-400 μl lysis buffer (1% Triton X-100, 50 mM Tris pH7.5, 300 mM NaCl, 5 mM EDTA, 1:1000 Protease inhibitor cocktail, 0.2 mM PMSF), and 3 wells were pooled for each IP, followed by centrifugation at 16,200 g 4°C for 10 minutes. For Fe^2+^ treatment, HEK cells were treated with 500 μM Fe^2+^ (Ammonium Iron(II) Sulfate Hexahydrate, 203505, Sigma-Aldrich) for 20-21 hours, and lysed in the same buffer where EDTA was replaced with 500 μM Fe^2+^. Supernatant was then incubated with 7 μl rabbit anti-MCU (26312-1-AP, ProteinTech), 7 μl rabbit IgG (2729S, cell signaling), or 5 μl rabbit anti-Flag (F7425, Sigma-Aldrich) for 2-3 hours at 4°C, and incubated with Protein A–Sepharose beads (17-0780-01, GE Healthcare) for another hour at 4°C. To detect iron levels, the colorimetric iron detection assay kit (ab83366, Abcam) was used according to the manufacturer’s instruction. Briefly, 180 μl Iron Assay Buffer was added to the beads with IP, and 100 μl of mixed buffer-beads was added to each well of total 2 wells. Next, 5 μl Iron Reducer was added to the well for iron (II+III), and 5 μl of Iron Assay Buffer was added to the well for iron (II), incubated at 37°C for 30 minutes. Then, 100 μl Iron Probe was added to each well, incubated at 37°C for 60 minutes, protected from light. Samples were measured immediately by an InfiniteM200 Pro TECAN plate reader at 593 nm.

#### ICP-MS

Iron concentration determinations were performed on an Agilent 8900 triple quadrupole inductively coupled plasma mass spectrometer (ICP-MS) attached to an Agilent SPS-4 autosampler. The sample introduction system also included a standard Scott double pass cooled spray chamber operated at 2’C, a 2.5mm i.d. Agilent glass torch, and a 200 μl/min Micromist nebulizer. The ICP-MS was fitted with the high sensitivity lens (s-lens), nickel cones and operated in ammonia (NH_3_) gas mode. Ammonia gas in the reaction cell of the triple quadrupole diminished the presence of argide including ArO which could form an isobaric interference with iron at mass 56. The large argide peak could also cause high backgrounds and more complex interferences at mass 57, another mass frequently monitored for iron concentrations. In ammonia gas mode, mass 56 and 57 backgrounds were dramatically reduced, resulting in an increase in the signal to noise ratio. This improved the iron detection limit in measured solutions on our instrument from about 1.8 ppb to 0.04 ppb. All reported concentrations were determined at mass 57 in ammonia gas mode. Before analysis, all solutions were further diluted by 50% by teeing in an internal standard solution of 1 μl/l indium (High Purity Standards) prepared with 2% v/v nitric acid to correct for instrumental drift. Procedural blanks were run in parallel with samples and averaged 48 pg contributing to less than 4% of the analytical signal and were regarded as negligible. Reproducibility was estimated from repeated analyses (n=3) of each sample in a single session.

#### PMBC Analysis

PBMCs were thawed in a 37°C water bath and plated into a 6-well plate (353046, Falcon) with RPMI media (11875-093, Gibco), supplemented with 10% fetal bovine serum (900-108, heat-inactivated, Gemini Bio-Products), 1% Pen-Strep (15140-122, Gibco), 1×GlutaMax (35050061, Thermo Fisher), and maintained in a 37°C, 5% CO_2_ incubator with humidified atmosphere. After a 1-hour recovery in the incubator, cells were then trypsinized with 0.25% Trypsin-EDTA (25200-056, Gibco) and collected in a cell suspension to be counted via TC20 Cell Counter (Bio-Rad).

Western blotting: After counting, the cell suspension (total 8 million cells) was equally split into 8 groups and replated with 3 ml media per well, 2 groups remaining untreated (to serve as controls), 3 groups treated with a combination of 20 μM Antimycin A (A8674, Sigma-Aldrich) and 2 μM Oligomycin (75351, Sigma-Aldrich) (AO), and 3 groups treated with 40 μM CCCP (C2759, Sigma-Aldrich). The untreated groups, serving as the 0-hour treatment in a time course, were then immediately pelleted and lysed with 150 μl lysis buffer (1% Triton X-100, 50 mM Tris, 300 mM NaCl, 5 mM EDTA, 0.2 mM PMSF, 1:1,000 Protease Inhibitor Cocktail) and stored at -80°C. At each of the following time points after treatment – 1 hour, 6 hours, and 14 hours – 1 group from each of the treatments (AO or CCCP) was similarly pelleted, lysed with lysis buffer, and stored at -80°C. Lysates collected were run in SDS-PAGE (30 μl loaded) and blotted with the following primary antibodies: mouse anti-Miro1 (WH0055288M1, Sigma-Aldrich) at 1:1,000, rabbit anti-GAPDH (5174S, Cell Signaling Technology) at 1:1,000, mouse anti-ATP5β (ab14730, Abcam) at 1:1,000, rabbit anti-VDAC (4661S, Cell Signaling Technology) at 1:1,000, and rabbit anti-β-Actin (4967S, Cell Signaling Technology) at 1:1,000.

ELISA: After counting, the cell suspension was equally split and replated with 3 ml media per well containing either 40 μM CCCP or a commensurate volume of DMSO for 6 hours. PBMC vials were provided in duplicate by MJFF, so the cell suspensions did not need to be split equally – they kept separate throughout the counting and replating process. For PBMCs that received drug treatment, the cell suspension was equally split into 3 wells. One well was pretreated with 10 μM Benidipine or MR3 (1915758622, MCULE) for 18 hours, and then this well and a second well were treated with 40 μM CCCP for 6 hours. The same volume of DMSO was applied to control wells. The minimum cell number needed for ELISA was 400,000 per well. Next, cells were scraped gently with a cell lifter and collected into suspension. The cells were then pelleted and lysed with lysis buffer (1% Triton X-100, 50 mM Tris, 300 mM NaCl, 5 mM EDTA, 0.2 mM PMSF, 1:1,000 Protease Inhibitor Cocktail) and stored at -80°C. The Miro1 protein level in each sample collected was determined using the RHOT1 ELISA Kit 96T (EKL54911, Biomatik) according to the manufacturer’s manual, and normalized to the total protein level in the same sample using the BCA Protein Assay Kit (23227, Thermo Scientific). For the ELISA kit, the specificity and stability were validated by Biomatik. The dynamic detection range (0.625-40 ng/ml), sensitivity (lower limit of detection–LLOD; 0.112 ng/ml), and precision (inter- and intra-assay) were determined by both Biomatik and us ^46^ (Figure S7), and the results were comparable. OD values for ELISA and BCA were measured on an InfiniteM200 Pro TECAN plate reader at 450 and 562 nm respectively, in duplicate. All experiments were performed in a double-blinded fashion, with PD status unknown to the experimenter.

Machine Learning: Anova analysis revealed a significant association of Miro1 Ratio with PD status, but not with other demographic and clinical parameters. We then trained a machine learning model to classify if a subject was with PD or not. The general form of the logistic regression model was represented by the following equation:

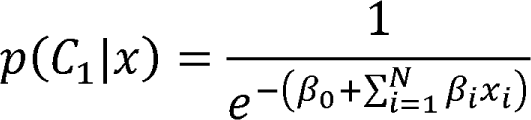

The goal of the model training was to learn the model parameters β_i_ such that we could predict the posterior probability *p*(*C_1_|x*). *C*_1_ represented the positive class “PD+” status and x represented the input of the model. We set a threshold value above which all predictions belonged to the positive class, i.e., “PD+”, and below which they belonged to the “PD-” class. We trained three logistic regression models with three different inputs. The first model used “Miro1 Ratio” as the only input, the second one had “UPDRS” as the only input, and the third model combined “Miro1 Ratio” and “UPDRS” as inputs. All three models were trained using the glm function in R. We broke the total subjects into a 60% training set and a 40% testing set. Data was cleaned by converting assigning column values to appropriate data types and omitting rows of data where no “Miro1 Ratio” or “UPDRS” values were present. The target variable, PD status, was labeled and encoded where 0 represented a subject without PD and 1 represented a PD positive subject. Table 1 at DOI: 10.17632/wtjcvm3cm3.2 showed the β_i_ and the corresponding p values. The estimate column represented β_i_ for each model. Since the “Miro1 Ratio” only and the “UPDRS” only models had a single feature, there were only two β_i_ values present. β_0_ represented the intercept and β_1_ represented “Miro1 Ratio” value or the “UPDRS” value depending on the model. We observed that the β_i_ estimate for the Miro1 Ratio model was significant (p<0.05) suggesting that the model predictions were statistically significant. Similarly, for UPDRS the β_1_ value was also significant. Both the UPDRS and Miro1 Ratio estimates were significant in the combined model, however the interaction term between them was non-significant. Once the models were trained, they were tested by evaluating the accuracy and AUC scores. Table 2 at DOI: 10.17632/wtjcvm3cm3.2 presented the accuracy and AUC scores for all models. We observed that the Miro1 Ratio only model performed worse with an accuracy of 67.6% than the UPDRS only model (81.2%) (Figure 1 at DOI: 10.17632/wtjcvm3cm3.2). However, together it yielded a higher score of 87.8%. We then added “Age” and “Sex” as additional inputs to the combined mode which did not affect the accuracy.

To determine the statistical significance between “Miro1 Ratio + UPDRS” and “UPDRS only”, we generated 100 samples (datasets) with replacement from each test set, predicted outcomes for each sample, and calculated the AUC scores for each sample. In line with standard bootstrapping sampling practice, we used the test set size as the sample size for generating each sample. Following the same process for both models it yielded two sample distributions, one of each model. Figure 2 at DOI: 10.17632/wtjcvm3cm3.2 showed the box plot for both distributions. More explanations and code are at DOI: 10.17632/wtjcvm3cm3.2.

#### Human Genetics

Based on our experimental data showing the importance of the calcium-Miro axis in PD models, the following calcium related genes were investigated for genetic association: *CACNA1S, CACNA1C, CACNA1D, CACNA1B, CACNA1G, CACNA1H, CACNA1I, RHOT1, RHOT2, MCU, CCDC109B, MICU1, MICU2, SMDT1, SLC8B1, TRPV1, TRPV2, TRPV3, TRPV4* (Table S5). First, to identify whether common variants regulating the expression (eQTL) or splicing (sQTL) of these genes are associated with PD risk, we queried the genome-wide association study (GWAS) summary statistics from ^47^. Genome-wide significant association (if any) was then intersected with eQTL/sQTL information from GTEx ^86^. Due to the absence of association, colocalization was not formally tested. Second, to test the association of rare variants in these genes we performed Burden and SKATO Test, as implemented in the R package *SKAT* v2.2.4, considering variants with a CADD (Combined Annotation Dependent Depletion) score ^87^ above 20 to only include variants most likely to have an impact on protein function. Additionally, we grouped variants in *CAC* genes by channel type, particularly grouping T-type (*CACNA1G, CACNA1H, CACNA1I*) and L-type (*CACNA1S, CACNA1C, CACNA1D*). For rare variant analysis, we obtained whole-genome sequencing (WGS) data from the Accelerating Medicines Partnership–Parkinson’s Disease (AMP–PD), composed of the following cohorts: New Discovery of Biomarkers (BioFIND), the Harvard Biomarker Study (HBS), the Parkinson’s Progression Markers Initiative (PPMI) ^88^, and the Parkinson’s Disease Biomarkers Program (PDBP). European ancestry individuals and principal components to account for genetic ancestry were computed as described in ^89^, leading to a total of 1,168 controls (45.4% males) and 2,241 cases (62.8% males). Significance was assessed separately for Burden and SKATO Test, though in genes where deleterious missense variants were all associated with the phenotype in the same direction (either increasing or decreasing risk) these two tests led to similar results.

#### Graphic Abstract

It was created with BioRender.com.

#### Quantification and statistical analysis

Throughout the paper, the distribution of data points is expressed as box-whisker or dot-plot with Mean±SEM, except otherwise stated. Box center line is median and box limits are upper and lower quartiles. One-Way or Two-Way ANOVA was performed for comparing multiple groups. Mann-Whitney *U* or T Test was performed for comparing two groups. Statistical analyses were performed using the Prism software (v8.01), Excel (v16.51), or R package *SKAT* (v2.2.4). For all experiments, between 3 and 107 cells, flies, humans, or independent experiments were used or performed. The number and meaning of experimental replications (n) can be found in corresponding Figure Legends. No statistical methods were used to predetermine sample sizes, but the number of flies, experiments, and biological replicates were chosen based on the nature of the experiments (it is usually difficult to assess an outcome that follows a normal distribution in our experiments), degree of variations, and published papers describing similar experiments. We did not exclude any data. Precise p values are labeled in figures.

## Supplementary Table Titles

**Table S1: Demographic Information of Postmortem Brain Donors, Related to** Figure 3.

**Table S2: Hit Compounds from the HTP Screening, Related to** Figure 6.

**Table S3: Pathway Analysis, Related to** Figure 6.

**Table S4: Demographic Information of PBMC Donors and Miro1 Values, Related to** Figure 7.

**Table S5: Human Genetic Analysis.**

## References

1. Apicco, D.J., Shlevkov, E., Nezich, C.L., Tran, D.T., Guilmette, E., Nicholatos, J.W., Bantle, C.M., Chen, Y., Glajch, K.E., Abraham, N.A., et al. (2021). The Parkinson’s disease-associated gene ITPKB protects against alpha-synuclein aggregation by regulating ER-to-mitochondria calcium release. Proceedings of the National Academy of Sciences of the United States of America 118. 10.1073/pnas.2006476118.

2. Lee, K.S., Huh, S., Lee, S., Wu, Z., Kim, A.K., Kang, H.Y., and Lu, B. (2018). Altered ER-mitochondria contact impacts mitochondria calcium homeostasis and contributes to neurodegeneration in vivo in disease models. Proceedings of the National Academy of Sciences of the United States of America 115, E8844–E8853. 10.1073/pnas.1721136115.

3. Tabata, Y., Imaizumi, Y., Sugawara, M., Andoh-Noda, T., Banno, S., Chai, M., Sone, T., Yamazaki, K., Ito, M., Tsukahara, K., et al. (2018). T-type Calcium Channels Determine the Vulnerability of Dopaminergic Neurons to Mitochondrial Stress in Familial Parkinson Disease. Stem cell reports 11, 1171–1184. 10.1016/j.stemcr.2018.09.006.

4. Buttner, S., Faes, L., Reichelt, W.N., Broeskamp, F., Habernig, L., Benke, S., Kourtis, N., Ruli, D., Carmona-Gutierrez, D., Eisenberg, T., et al. (2013). The Ca2+/Mn2+ ion-pump PMR1 links elevation of cytosolic Ca(2+) levels to alpha-synuclein toxicity in Parkinson’s disease models. Cell death and differentiation 20, 465–477. 10.1038/cdd.2012.142.

5. Surmeier, D.J., Obeso, J.A., and Halliday, G.M. (2017). Selective neuronal vulnerability in Parkinson disease. Nature reviews. Neuroscience 18, 101–113. 10.1038/nrn.2016.178.

6. Kim, J.W., Yin, X., Jhaldiyal, A., Khan, M.R., Martin, I., Xie, Z., Perez-Rosello, T., Kumar, M., Abalde-Atristain, L., Xu, J., et al. (2020). Defects in mRNA Translation in LRRK2-Mutant hiPSC-Derived Dopaminergic Neurons Lead to Dysregulated Calcium Homeostasis. Cell stem cell 27, 633–645 e637. 10.1016/j.stem.2020.08.002.

7. Angelova, P.R., Choi, M.L., Berezhnov, A.V., Horrocks, M.H., Hughes, C.D., De, S., Rodrigues, M., Yapom, R., Little, D., Dolt, K.S., et al. (2020). Alpha synuclein aggregation drives ferroptosis: an interplay of iron, calcium and lipid peroxidation. Cell death and differentiation 27, 2781–2796. 10.1038/s41418-020-0542-z.

8. Verma, M., Callio, J., Otero, P.A., Sekler, I., Wills, Z.P., and Chu, C.T. (2017). Mitochondrial Calcium Dysregulation Contributes to Dendrite Degeneration Mediated by PD/LBD-Associated LRRK2 Mutants. The Journal of neuroscience : the official journal of the Society for Neuroscience 37, 11151–11165. 10.1523/JNEUROSCI.3791-16.2017.

9. Belaidi, A.A., and Bush, A.I. (2016). Iron neurochemistry in Alzheimer’s disease and Parkinson’s disease: targets for therapeutics. Journal of neurochemistry 139 *Suppl 1*, 179–197. 10.1111/jnc.13425.

10. Vuuren, M.J.V., Nell, T.A., Carr, J.A., Kell, D.B., and Pretorius, E. (2020). Iron Dysregulation and Inflammagens Related to Oral and Gut Health Are Central to the Development of Parkinson’s Disease. Biomolecules 11. 10.3390/biom11010030.

11. Baughman, J.M., Perocchi, F., Girgis, H.S., Plovanich, M., Belcher-Timme, C.A., Sancak, Y., Bao, X.R., Strittmatter, L., Goldberger, O., Bogorad, R.L., et al. (2011). Integrative genomics identifies MCU as an essential component of the mitochondrial calcium uniporter. Nature 476, 341–345. 10.1038/nature10234.

12. Palty, R., Silverman, W.F., Hershfinkel, M., Caporale, T., Sensi, S.L., Parnis, J., Nolte, C., Fishman, D., Shoshan-Barmatz, V., Herrmann, S., et al. (2010). NCLX is an essential component of mitochondrial Na+/Ca2+ exchange. Proceedings of the National Academy of Sciences of the United States of America 107, 436–441. 10.1073/pnas.0908099107.

13. Fan, C., Fan, M., Orlando, B.J., Fastman, N.M., Zhang, J., Xu, Y., Chambers, M.G., Xu, X., Perry, K., Liao, M., and Feng, L. (2018). X-ray and cryo-EM structures of the mitochondrial calcium uniporter. Nature 559, 575–579. 10.1038/s41586-018-0330-9.

14. Fan, M., Zhang, J., Tsai, C.W., Orlando, B.J., Rodriguez, M., Xu, Y., Liao, M., Tsai, M.F., and Feng, L. (2020). Structure and mechanism of the mitochondrial Ca(2+) uniporter holocomplex. Nature 582, 129–133. 10.1038/s41586-020-2309-6.

15. Lambert, J.P., Luongo, T.S., Tomar, D., Jadiya, P., Gao, E., Zhang, X., Lucchese, A.M., Kolmetzky, D.W., Shah, N.S., and Elrod, J.W. (2019). MCUB Regulates the Molecular Composition of the Mitochondrial Calcium Uniporter Channel to Limit Mitochondrial Calcium Overload During Stress. Circulation 140, 1720–1733. 10.1161/CIRCULATIONAHA.118.037968.

16. Patron, M., Tarasenko, D., Nolte, H., Kroczek, L., Ghosh, M., Ohba, Y., Lasarzewski, Y., Ahmadi, Z.A., Cabrera-Orefice, A., Eyiama, A., et al. (2022). Regulation of mitochondrial proteostasis by the proton gradient. The EMBO journal 41, e110476. 10.15252/embj.2021110476.

17. Paul, B.T., Manz, D.H., Torti, F.M., and Torti, S.V. (2017). Mitochondria and Iron: current questions. Expert Rev Hematol 10, 65–79. 10.1080/17474086.2016.1268047.

18. Dong, Z., Shanmughapriya, S., Tomar, D., Siddiqui, N., Lynch, S., Nemani, N., Breves, S.L., Zhang, X., Tripathi, A., Palaniappan, P., et al. (2017). Mitochondrial Ca(2+) Uniporter Is a Mitochondrial Luminal Redox Sensor that Augments MCU Channel Activity. Molecular cell 65, 1014–1028 e1017. 10.1016/j.molcel.2017.01.032.

19. Berridge, M.J., Bootman, M.D., and Roderick, H.L. (2003). Calcium signalling: dynamics, homeostasis and remodelling. Nature reviews. Molecular cell biology 4, 517–529. 10.1038/nrm1155.

20. Clapham, D.E. (2007). Calcium signaling. Cell 131, 1047–1058. 10.1016/j.cell.2007.11.028.

21. Tomar, D., Dong, Z., Shanmughapriya, S., Koch, D.A., Thomas, T., Hoffman, N.E., Timbalia, S.A., Goldman, S.J., Breves, S.L., Corbally, D.P., et al. (2016). MCUR1 Is a Scaffold Factor for the MCU Complex Function and Promotes Mitochondrial Bioenergetics. Cell Rep 15, 1673–1685. 10.1016/j.celrep.2016.04.050.

22. Ghosh, S., Basu Ball, W., Madaris, T.R., Srikantan, S., Madesh, M., Mootha, V.K., and Gohil, V.M. (2020). An essential role for cardiolipin in the stability and function of the mitochondrial calcium uniporter. Proceedings of the National Academy of Sciences of the United States of America 117, 16383–16390. 10.1073/pnas.2000640117.

23. Michailides, C., and Velissaris, D. (2021). Common anti-oxidant vitamin C as an anti-infective agent with remedial role on SARS-CoV-2 infection. An update. Monaldi Arch Chest Dis 91. 10.4081/monaldi.2021.1808.

24. Chen, A.Y., Lu, J.M., Yao, Q., and Chen, C. (2016). Entacapone is an Antioxidant More Potent than Vitamin C and Vitamin E for Scavenging of Hypochlorous Acid and Peroxynitrite, and the Inhibition of Oxidative Stress-Induced Cell Death. Med Sci Monit 22, 687–696. 10.12659/msm.896462.

25. Munson, M.J., Mathai, B.J., Ng, M.Y.W., Trachsel-Moncho, L., de la Ballina, L.R., Schultz, S.W., Aman, Y., Lystad, A.H., Singh, S., Singh, S., et al. (2021). GAK and PRKCD are positive regulators of PRKN-independent mitophagy. Nature communications 12, 6101. 10.1038/s41467-021-26331-7.

26. Billesbolle, C.B., Azumaya, C.M., Kretsch, R.C., Powers, A.S., Gonen, S., Schneider, S., Arvedson, T., Dror, R.O., Cheng, Y., and Manglik, A. (2020). Structure of hepcidin-bound ferroportin reveals iron homeostatic mechanisms. Nature 586, 807–811. 10.1038/s41586-020-2668-z.

27. Lin, Y.F., Cheng, C.W., Shih, C.S., Hwang, J.K., Yu, C.S., and Lu, C.H. (2016). MIB: Metal Ion-Binding Site Prediction and Docking Server. J Chem Inf Model 56, 2287–2291. 10.1021/acs.jcim.6b00407.

28. Lu, C.H., Lin, Y.F., Lin, J.J., and Yu, C.S. (2012). Prediction of metal ion-binding sites in proteins using the fragment transformation method. PloS one 7, e39252. 10.1371/journal.pone.0039252.

29. Lee, S.K., Shanmughapriya, S., Mok, M.C.Y., Dong, Z., Tomar, D., Carvalho, E., Rajan, S., Junop, M.S., Madesh, M., and Stathopulos, P.B. (2016). Structural Insights into Mitochondrial Calcium Uniporter Regulation by Divalent Cations. Cell Chem Biol 23, 1157–1169. 10.1016/j.chembiol.2016.07.012.

30. Fernandez, B., Olmedo, P., Gil, F., Fdez, E., Naaldijk, Y., Rivero-Rios, P., Bracher, F., Grimm, C., Churchill, G.C., and Hilfiker, S. (2022). Iron-induced cytotoxicity mediated by endolysosomal TRPML1 channels is reverted by TFEB. Cell Death Dis 13, 1047. 10.1038/s41419-022-05504-2.

31. Jadiya, P., Kolmetzky, D.W., Tomar, D., Di Meco, A., Lombardi, A.A., Lambert, J.P., Luongo, T.S., Ludtmann, M.H., Pratico, D., and Elrod, J.W. (2019). Impaired mitochondrial calcium efflux contributes to disease progression in models of Alzheimer’s disease. Nature communications 10, 3885. 10.1038/s41467-019-11813-6.

32. Hsieh, C.H., Shaltouki, A., Gonzalez, A.E., Bettencourt da Cruz, A., Burbulla, L.F., St Lawrence, E., Schule, B., Krainc, D., Palmer, T.D., and Wang, X. (2016). Functional Impairment in Miro Degradation and Mitophagy Is a Shared Feature in Familial and Sporadic Parkinson’s Disease. Cell stem cell 19, 709–724. 10.1016/j.stem.2016.08.002.

33. Shaltouki, A., Hsieh, C.H., Kim, M.J., and Wang, X. (2018). Alpha-synuclein delays mitophagy and targeting Miro rescues neuron loss in Parkinson’s models. Acta Neuropathol 136, 607–620. 10.1007/s00401-018-1873-4.

34. Hsieh, C.H., Li, L., Vanhauwaert, R., Nguyen, K.T., Davis, M.D., Bu, G., Wszolek, Z.K., and Wang, X. (2019). Miro1 Marks Parkinson’s Disease Subset and Miro1 Reducer Rescues Neuron Loss in Parkinson’s Models. Cell metabolism 30, 1131–1140 e1137. 10.1016/j.cmet.2019.08.023.

35. Li, L., Conradson, D.M., Bharat, V., Kim, M.J., Hsieh, C.H., Minhas, P.S., Papakyrikos, A.M., Durairaj, A.S., Ludlam, A., Andreasson, K.I., et al. (2021). A mitochondrial membrane-bridging machinery mediates signal transduction of intramitochondrial oxidation. Nat Metab. 10.1038/s42255-021-00443-2.

36. Lee, D.G., Park, J., Lee, H.S., Lee, S.R., and Lee, D.S. (2016). Iron overload-induced calcium signals modulate mitochondrial fragmentation in HT-22 hippocampal neuron cells. Toxicology 365, 17–24. 10.1016/j.tox.2016.07.022.

37. Niescier, R.F., Hong, K., Park, D., and Min, K.T. (2018). MCU Interacts with Miro1 to Modulate Mitochondrial Functions in Neurons. The Journal of neuroscience : the official journal of the Society for Neuroscience 38, 4666–4677. 10.1523/JNEUROSCI.0504-18.2018.

38. Gottschalk, B., Klec, C., Leitinger, G., Bernhart, E., Rost, R., Bischof, H., Madreiter-Sokolowski, C.T., Radulovic, S., Eroglu, E., Sattler, W., et al. (2019). MICU1 controls cristae junction and spatially anchors mitochondrial Ca(2+) uniporter complex. Nature communications 10, 3732. 10.1038/s41467-019-11692-x.

39. Katona, M., Bartok, A., Nichtova, Z., Csordas, G., Berezhnaya, E., Weaver, D., Ghosh, A., Varnai, P., Yule, D.I., and Hajnoczky, G. (2022). Capture at the ER-mitochondrial contacts licenses IP(3) receptors to stimulate local Ca(2+) transfer and oxidative metabolism. Nature communications 13, 6779. 10.1038/s41467-022-34365-8.

40. Garcia-Rivas Gde, J., Carvajal, K., Correa, F., and Zazueta, C. (2006). Ru360, a specific mitochondrial calcium uptake inhibitor, improves cardiac post-ischaemic functional recovery in rats in vivo. Br J Pharmacol 149, 829–837. 10.1038/sj.bjp.0706932.

41. Wang, X., Winter, D., Ashrafi, G., Schlehe, J., Wong, Y.L., Selkoe, D., Rice, S., Steen, J., LaVoie, M.J., and Schwarz, T.L. (2011). PINK1 and Parkin target Miro for phosphorylation and degradation to arrest mitochondrial motility. Cell 147, 893–906. 10.1016/j.cell.2011.10.018.

42. Lopez-Domenech, G., Howden, J.H., Covill-Cooke, C., Morfill, C., Patel, J.V., Burli, R., Crowther, D., Birsa, N., Brandon, N.J., and Kittler, J.T. (2021). Loss of neuronal Miro1 disrupts mitophagy and induces hyperactivation of the integrated stress response. The EMBO journal 40, e100715. 10.15252/embj.2018100715.

43. Wang, X., and Schwarz, T.L. (2009). The mechanism of Ca2+ -dependent regulation of kinesin-mediated mitochondrial motility. Cell 136, 163–174. 10.1016/j.cell.2008.11.046.

44. Darakhshan, S., and Pour, A.B. (2015). Tranilast: a review of its therapeutic applications. Pharmacol Res 91, 15–28. 10.1016/j.phrs.2014.10.009.

45. Ordureau, A., Paulo, J.A., Zhang, J., An, H., Swatek, K.N., Cannon, J.R., Wan, Q., Komander, D., and Harper, J.W. (2020). Global Landscape and Dynamics of Parkin and USP30-Dependent Ubiquitylomes in iNeurons during Mitophagic Signaling. Molecular cell 77, 1124–1142 e1110. 10.1016/j.molcel.2019.11.013.

46. Nguyen, D., Bharat, V., Conradson, D.M., Nandakishore, P., and Wang, X. (2021). Miro1 Impairment in a Parkinson’s At-Risk Cohort. Front Mol Neurosci 14, 734273. 10.3389/fnmol.2021.734273.

47. Nalls, M.A., Blauwendraat, C., Vallerga, C.L., Heilbron, K., Bandres-Ciga, S., Chang, D., Tan, M., Kia, D.A., Noyce, A.J., Xue, A., et al. (2019). Identification of novel risk loci, causal insights, and heritable risk for Parkinson’s disease: a meta-analysis of genome-wide association studies. The Lancet. Neurology 18, 1091–1102. 10.1016/S1474-4422(19)30320-5.

48. Papapetropoulos, S., Lee, M.S., Boyer, S., and Newbold, E.J. (2019). A Phase 2, Randomized, Double-Blind, Placebo-Controlled Trial of CX-8998, a Selective Modulator of the T-Type Calcium Channel in Inadequately Treated Moderate to Severe Essential Tremor: T-CALM Study Design and Methodology for Efficacy Endpoint and Digital Biomarker Selection. Front Neurol 10, 597. 10.3389/fneur.2019.00597.

49. Margaret S. Lee, S.P., Michelle S. Higgin, Muralikrishna Duvvuri, Bruce Rehlaender, Evan NEWBOLD (2020) Treating essential tremor using (r)-2-(4-isopropylphenyl)-n-(1-(5-(2,2,2-trifluoroethoxy)pyridin-2-yl)ethyl)acetamide. patent WO2020072773A1, 2020-04-09.

50. Xiang, Z., Thompson, A.D., Brogan, J.T., Schulte, M.L., Melancon, B.J., Mi, D., Lewis, L.M., Zou, B., Yang, L., Morrison, R., et al. (2011). The Discovery and Characterization of ML218: A Novel, Centrally Active T-Type Calcium Channel Inhibitor with Robust Effects in STN Neurons and in a Rodent Model of Parkinson’s Disease. ACS Chem Neurosci 2, 730–742. 10.1021/cn200090z.

51. Parkinson Study Group, S.-P.D.I.I.I.I. (2020). Isradipine Versus Placebo in Early Parkinson Disease: A Randomized Trial. Ann Intern Med 172, 591–598. 10.7326/M19-2534.

52. Hess, C.W., and Okun, M.S. (2016). Diagnosing Parkinson Disease. Continuum (Minneap Minn) 22, 1047–1063. 10.1212/CON.0000000000000345.

53. Bartels, T. (2016). Conformation-Specific Detection of alpha-Synuclein: The Search for a Biomarker in Parkinson Disease. JAMA Neurol. 10.1001/jamaneurol.2016.4813.

54. Bharat, V., and Wang, X. (2020). Precision Neurology for Parkinson’s Disease: Coupling Miro1-Based Diagnosis With Drug Discovery. Movement disorders : official journal of the Movement Disorder Society 35, 1502–1508. 10.1002/mds.28194.

55. Choi, M.L., Chappard, A., Singh, B.P., Maclachlan, C., Rodrigues, M., Fedotova, E.I., Berezhnov, A.V., De, S., Peddie, C.J., Athauda, D., et al. (2022). Pathological structural conversion of alpha-synuclein at the mitochondria induces neuronal toxicity. Nature neuroscience 25, 1134–1148. 10.1038/s41593-022-01140-3.

56. Ramezani, M., Wagenknecht-Wiesner, A., Wang, T., Holowka, D.A., Eliezer, D., and Baird, B.A. (2023). Alpha Synuclein Modulates Mitochondrial Ca (2+) Uptake from ER During Cell Stimulation and Under Stress Conditions. bioRxiv. 10.1101/2023.04.23.537965.

57. Erustes, A.G., Guarache, G.C., Guedes, E.D.C., Leao, A., Pereira, G., and Smaili, S.S. (2022). alpha-Synuclein Interactions in Mitochondria-ER Contacts: A Possible Role in Parkinson’s Disease. Contact (Thousand Oaks) 5, 25152564221119347. 10.1177/25152564221119347.

58. Melo, T.Q., Copray, S., and Ferrari, M.F.R. (2018). Alpha-Synuclein Toxicity on Protein Quality Control, Mitochondria and Endoplasmic Reticulum. Neurochem Res 43, 2212–2223. 10.1007/s11064-018-2673-x.

59. Paillusson, S., Gomez-Suaga, P., Stoica, R., Little, D., Gissen, P., Devine, M.J., Noble, W., Hanger, D.P., and Miller, C.C.J. (2017). alpha-Synuclein binds to the ER-mitochondria tethering protein VAPB to disrupt Ca(2+) homeostasis and mitochondrial ATP production. Acta Neuropathol 134, 129–149. 10.1007/s00401-017-1704-z.

60. Devi, L., Raghavendran, V., Prabhu, B.M., Avadhani, N.G., and Anandatheerthavarada, H.K. (2008). Mitochondrial import and accumulation of alpha-synuclein impair complex I in human dopaminergic neuronal cultures and Parkinson disease brain. The Journal of biological chemistry 283, 9089–9100. 10.1074/jbc.M710012200.

61. Kamer, K.J., Sancak, Y., Fomina, Y., Meisel, J.D., Chaudhuri, D., Grabarek, Z., and Mootha, V.K. (2018). MICU1 imparts the mitochondrial uniporter with the ability to discriminate between Ca(2+) and Mn(2+). Proceedings of the National Academy of Sciences of the United States of America 115, E7960–E7969. 10.1073/pnas.1807811115.

62. Chan, N.C., Salazar, A.M., Pham, A.H., Sweredoski, M.J., Kolawa, N.J., Graham, R.L., Hess, S., and Chan, D.C. (2011). Broad activation of the ubiquitin-proteasome system by Parkin is critical for mitophagy. Human molecular genetics 20, 1726–1737. 10.1093/hmg/ddr048.

63. Konig, T., Nolte, H., Aaltonen, M.J., Tatsuta, T., Krols, M., Stroh, T., Langer, T., and McBride, H.M. (2021). MIROs and DRP1 drive mitochondrial-derived vesicle biogenesis and promote quality control. Nature cell biology. 10.1038/s41556-021-00798-4.

64. Melentijevic, I., Toth, M.L., Arnold, M.L., Guasp, R.J., Harinath, G., Nguyen, K.C., Taub, D., Parker, J.A., Neri, C., Gabel, C.V., et al. (2017). C. elegans neurons jettison protein aggregates and mitochondria under neurotoxic stress. Nature 542, 367–371. 10.1038/nature21362.

65. Davis, C.H., Kim, K.Y., Bushong, E.A., Mills, E.A., Boassa, D., Shih, T., Kinebuchi, M., Phan, S., Zhou, Y., Bihlmeyer, N.A., et al. (2014). Transcellular degradation of axonal mitochondria. Proceedings of the National Academy of Sciences of the United States of America 111, 9633–9638. 10.1073/pnas.1404651111.

66. Saha, T., Dash, C., Jayabalan, R., Khiste, S., Kulkarni, A., Kurmi, K., Mondal, J., Majumder, P.K., Bardia, A., Jang, H.L., and Sengupta, S. (2022). Intercellular nanotubes mediate mitochondrial trafficking between cancer and immune cells. Nat Nanotechnol 17, 98–106. 10.1038/s41565-021-01000-4.

67. Rosina, M., Ceci, V., Turchi, R., Chuan, L., Borcherding, N., Sciarretta, F., Sanchez-Diaz, M., Tortolici, F., Karlinsey, K., Chiurchiu, V., et al. (2022). Ejection of damaged mitochondria and their removal by macrophages ensure efficient thermogenesis in brown adipose tissue. Cell Metab. 10.1016/j.cmet.2022.02.016.

68. Ahmad, T., Mukherjee, S., Pattnaik, B., Kumar, M., Singh, S., Kumar, M., Rehman, R., Tiwari, B.K., Jha, K.A., Barhanpurkar, A.P., et al. (2014). Miro1 regulates intercellular mitochondrial transport & enhances mesenchymal stem cell rescue efficacy. The EMBO journal 33, 994–1010. 10.1002/embj.201386030.

69. Trinh, K., Moore, K., Wes, P.D., Muchowski, P.J., Dey, J., Andrews, L., and Pallanck, L.J. (2008). Induction of the phase II detoxification pathway suppresses neuron loss in Drosophila models of Parkinson’s disease. The Journal of neuroscience : the official journal of the Society for Neuroscience 28, 465–472. 10.1523/JNEUROSCI.4778-07.2008.

70. Tsai, P.I., Course, M.M., Lovas, J.R., Hsieh, C.H., Babic, M., Zinsmaier, K.E., and Wang, X. (2014). PINK1-mediated phosphorylation of Miro inhibits synaptic growth and protects dopaminergic neurons in Drosophila. Scientific reports 4, 6962. 10.1038/srep06962.

71. Castillo-Quan, J.I., Li, L., Kinghorn, K.J., Ivanov, D.K., Tain, L.S., Slack, C., Kerr, F., Nespital, T., Thornton, J., Hardy, J., et al. (2016). Lithium Promotes Longevity through GSK3/NRF2-Dependent Hormesis. Cell Rep 15, 638–650. 10.1016/j.celrep.2016.03.041.

72. Sofola, O., Kerr, F., Rogers, I., Killick, R., Augustin, H., Gandy, C., Allen, M.J., Hardy, J., Lovestone, S., and Partridge, L. (2010). Inhibition of GSK-3 ameliorates Abeta pathology in an adult-onset Drosophila model of Alzheimer’s disease. PLoS genetics 6, e1001087. 10.1371/journal.pgen.1001087.

73. Byers, B., Cord, B., Nguyen, H.N., Schule, B., Fenno, L., Lee, P.C., Deisseroth, K., Langston, J.W., Pera, R.R., and Palmer, T.D. (2011). SNCA triplication Parkinson’s patient’s iPSC-derived DA neurons accumulate alpha-synuclein and are susceptible to oxidative stress. PloS one 6, e26159. 10.1371/journal.pone.0026159.

74. Kriks, S., Shim, J.W., Piao, J., Ganat, Y.M., Wakeman, D.R., Xie, Z., Carrillo-Reid, L., Auyeung, G., Antonacci, C., Buch, A., et al. (2011). Dopamine neurons derived from human ES cells efficiently engraft in animal models of Parkinson’s disease. Nature 480, 547–551. 10.1038/nature10648.

75. Wichterle, H., Lieberam, I., Porter, J.A., and Jessell, T.M. (2002). Directed differentiation of embryonic stem cells into motor neurons. Cell 110, 385–397.

76. Nguyen, H.N., Byers, B., Cord, B., Shcheglovitov, A., Byrne, J., Gujar, P., Kee, K., Schule, B., Dolmetsch, R.E., Langston, W., et al. (2011). LRRK2 mutant iPSC-derived DA neurons demonstrate increased susceptibility to oxidative stress. Cell stem cell 8, 267–280. 10.1016/j.stem.2011.01.013.

77. Sanders, L.H., Laganiere, J., Cooper, O., Mak, S.K., Vu, B.J., Huang, Y.A., Paschon, D.E., Vangipuram, M., Sundararajan, R., Urnov, F.D., et al. (2014). LRRK2 mutations cause mitochondrial DNA damage in iPSC-derived neural cells from Parkinson’s disease patients: reversal by gene correction. Neurobiology of disease 62, 381–386. 10.1016/j.nbd.2013.10.013.

78. Birsa, N., Norkett, R., Wauer, T., Mevissen, T.E., Wu, H.C., Foltynie, T., Bhatia, K., Hirst, W.D., Komander, D., Plun-Favreau, H., and Kittler, J.T. (2014). Lysine 27 ubiquitination of the mitochondrial transport protein Miro is dependent on serine 65 of the Parkin ubiquitin ligase. The Journal of biological chemistry 289, 14569–14582. 10.1074/jbc.M114.563031.

79. Hiroyuki Katayama, H.H., Koji Nagasawa, Hiroshi Kurokawa, Mayu Sugiyama, Ryoko Ando, Masaaki Funata, Nobuyo Yoshida, Misaki Homma, Takanori Nishimura, Megumu Takahashi, Yoko Ishida, Hiroyuki Hioki, Yoshiyuki Tsujihata, and Atsushi Miyawaki (2020). Visualizing and Modulating Mitophagy for Therapeutic Studies of Neurodegeneration. Cell 181, 12. 10.1016/j.cell.2020.04.025.

80. Hoshino, A., Wang, W.J., Wada, S., McDermott-Roe, C., Evans, C.S., Gosis, B., Morley, M.P., Rathi, K.S., Li, J., Li, K., et al. (2019). The ADP/ATP translocase drives mitophagy independent of nucleotide exchange. Nature 575, 375–379. 10.1038/s41586-019-1667-4.

81. Clark, E.H., Vazquez de la Torre, A., Hoshikawa, T., and Briston, T. (2020). Targeting mitophagy in Parkinson’s disease. The Journal of biological chemistry. 10.1074/jbc.REV120.014294.

82. Ehrenkaufer, G.M., Suresh, S., Solow-Cordero, D., and Singh, U. (2018). High-Throughput Screening of Entamoeba Identifies Compounds Which Target Both Life Cycle Stages and Which Are Effective Against Metronidazole Resistant Parasites. Front Cell Infect Microbiol 8, 276. 10.3389/fcimb.2018.00276.

83. Percha, B., and Altman, R.B. (2018). A global network of biomedical relationships derived from text. Bioinformatics 34, 2614–2624. 10.1093/bioinformatics/bty114.

84. Rival, T., Soustelle, L., Strambi, C., Besson, M.T., Iche, M., and Birman, S. (2004). Decreasing glutamate buffering capacity triggers oxidative stress and neuropil degeneration in the Drosophila brain. Current biology : CB 14, 599–605. 10.1016/j.cub.2004.03.039.

85. Kinghorn, K.J., Castillo-Quan, J.I., Bartolome, F., Angelova, P.R., Li, L., Pope, S., Cocheme, H.M., Khan, S., Asghari, S., Bhatia, K.P., et al. (2015). Loss of PLA2G6 leads to elevated mitochondrial lipid peroxidation and mitochondrial dysfunction. Brain : a journal of neurology 138, 1801–1816. 10.1093/brain/awv132.

86. Consortium, G.T., Laboratory, D.A., Coordinating Center -Analysis Working, G., Statistical Methods groups-Analysis Working, G., Enhancing, G.g., Fund, N.I.H.C., Nih/Nci, Nih/Nhgri, Nih/Nimh, Nih/Nida, et al. (2017). Genetic effects on gene expression across human tissues. Nature 550, 204–213. 10.1038/nature24277.

87. Rentzsch, P., Witten, D., Cooper, G.M., Shendure, J., and Kircher, M. (2019). CADD: predicting the deleteriousness of variants throughout the human genome. Nucleic Acids Res 47, D886–D894. 10.1093/nar/gky1016.

88. Nalls, M.A., Keller, M.F., Hernandez, D.G., Chen, L., Stone, D.J., Singleton, A.B., and Parkinson’s Progression Marker Initiative, i. (2016). Baseline genetic associations in the Parkinson’s Progression Markers Initiative (PPMI). Movement disorders : official journal of the Movement Disorder Society 31, 79–85. 10.1002/mds.26374.

89. Le Guen, Y., Napolioni, V., Belloy, M.E., Yu, E., Krohn, L., Ruskey, J.A., Gan-Or, Z., Kennedy, G., Eger, S.J., and Greicius, M.D. (2021). Common X-Chromosome Variants Are Associated with Parkinson Disease Risk. Ann Neurol 90, 22–34. 10.1002/ana.26051.

