## Supplementary material for "A Mitochondrial Inside-Out Iron-Calcium Signal Reveals Drug Targets for Parkinson’s Disease": Supplmentary Figures

#### Supplementary Figure Legend

**Figure S1. Cell Death Assay, SEC Blots, and ROS Assays.** (A) HEK cells were treated as indicated for 24 hours and cell viability was measured by the MTT assay. n=3. Not significant compared to "untreated". Two-tailed Welch's T Test. (B) HEK cells were treated as indicated and run through SEC. Each fraction containing differential MW of protein complexes was run in SDS-PAGE and blotted with anti-MCU. (C) Validation of anti-MCU using the SEC sample from wild-type HEK cells (control; *MCU+/+*) and lysate of *MCU-/-* HEK cells in SDS-PAGE. (D-E) Similar to (B) and Figure 1G. (D) MCU SEC profile of HEK cells treated with $H_2O_2$. (E) Miro1 SEC profiles. Note that $Fe^{2+}$ or $H_2O_2$ treatment shifted the elution peak fractions of MCU from fractions "14-18" to "4-8", whereas Miro1's peak fractions remained in fractions "10-12" under all conditions. (F) Similar to Figure 1C-D, mitochondrial $Ca^{2+}$ dynamics was measured. n=15 cells from 3 coverslips. (G-H) Similar to Figure 1E, MCU oligomerization was measured. n=4. (F-G) 500 μM $Fe^{2+}$ was added for 24 hours. 100 μM VitC or DFP was added for 16 hours. One-Way Anova Post Hoc Tukey Test. (H) 100 μM $H_2O_2$ was added for 30 minutes. Two-tailed Welch's T Test. Compared within each genotype. The same volume of the solvent was added to controls in all panels. See also Figure 1.

**Figure S2. Mitochondrial Protein Responses to Iron.** HEK cells were treated as indicated, lysates were run in SDS-PAGE, and blotted. The band intensity was normalized to that of GAPDH from the same blot. n=5-19 independent experiments. Precise p and n are in the figure. One-Way Anova Post Hoc Tukey Test. Condition 1 contained EDTA. For condition 3: 5 mM $Fe^{2+}$ was added to the media 20-22 hours before cells were lysed.

**Figure S3. PD Postmortem Brains and Validation of Models.** (A) Postmortem brains were run in Native- or SDS-PAGE and blotted. The band intensity normalized to the total protein level measured by BCA is divided by that of the universal control (CVD, cardiovascular disease) on the same blot, which was included on every blot. The MCU oligomer bands in Native-PAGE and the NCLX and MCUb bands in SDS-PAGE (average of 3 replicates) are used in the plot here and in Figure 3A. HC: healthy control. (B) Similar to Figure 3C and E, Fmax/F0 of the Rhod-2 intensity (left) and the inverted RPA/MitoTracker intensity was measured in neurons day 40 after differentiation. n=20 cell bodies from 3 independent coverslips. Two-tailed Welch's T Test. (C) iPSC-derived neurons treated as indicated, were immunostained with TUNEL and Dapi, and imaged under a confocal microscope. Scale bar: 50 μm. Below: Quantification of the percentage of TUNEL-positive neurons. n=20 images from 3 independent coverslips. Comparison within each genotype (P values are significant compared to every other condition; non-significant P not shown, One-Way Anova Post Hoc Tukey Test.). (D) Similar to Figure 4A, iPSC-derived neurons with or without treatment of 100 μM DFP for 24 hours were stimulated with thrombin, and mitochondrial $Ca^{2+}$ (Rhod-2) was measured. Quantification of the peak fluorescent intensity normalized to the baseline within each cell body. (E) Similar to Figure 3E, the inverted RPA/MitoTracker intensity was measured. (D-E) n=15 cell bodies from 3 independent coverslips. Compared to "HC-1". One-Way Anova Post Hoc Dunnett's Test. (F) In iPSC-derived neurons transfected with miRFP670_P2A_GCaMP6f-Miro1-Anchor and treated as indicated, the intensity of GCaMP-Miro1-Anchor is normalized to miRFP670 within the same cell body. n=15 cell bodies from 3 coverslips. One-Way Anova Post Hoc Tukey Test. Data without treatment is the same as in Figure 4G. (G) Flies with indicated genotypes were lysed and run in SDS-PAGE. n=4 independent experiments. Two-tailed Mann-Whitney Test. See also Figure 3-5.

**Figure S4. HTP Screens Identify Hit Compounds That Reduce Miro1 upon Depolarization.** Miro1 values are modulated by compounds from 3 libraries. The calculation of Relative Miro1 Intensity is described in Methods. Negative values indicate Miro1 reduction. Each black or red dot represents the Miro1 intensity value from ICC on the sporadic PD cell line treated with each unique compound combined with FCCP. Each compound was applied 10 hours before application of 20 μM FCCP for another 14 hours. Blue dots represent the Miro1 values of the same sporadic PD cell line treated only with FCCP for 14 hours from multiple independent experiments. Red dashed lines represent 3SD. (A) NIH Clinical Collection (screening concentration=10 μM, n=4 biological repeats). Right: Eleven primary hits; mean±SEM are shown. (B) Biomol FDA (5-fold screening concentrations, n=2 biological repeats; data of 20 μM are shown). Right: Three primary hits (purple, red, gray line) in a dose-dependent manner. The green line shows data from a compound that increases Miro1 levels, and the black line shows data from a non-responsive compound. $IC_{50}$: Allopurinol: 1.338 μM; Fenbufen: 6.260 μM; Astemizole: 12.63 μM. (C) Sigma LOPAC (screening concentration=20 μM, n=1 biological repeat). Right: Twenty-one primary hits. See also Figure 6.

**Figure S5. Independent Validation of Primary Hits.** (A) Representative images of a healthy (upper row) or a sporadic PD fibroblast cell line (middle and lower rows), treated as indicated, and stained with anti-Miro1 and Dapi. Scale bar: 100 μm. (B) Heat map shows the mean Miro1 protein intensities calculated from images similar to (A). Each mean Miro1 value from the sporadic PD line treated with a unique compound is expressed as a percentage of the mean value from the same PD line with no treatment (DMSO; 100). Green shows Miro1 reduction and magenta shows Miro1 elevation. n=4 (drug treatment), 16 (PD, FCCP), and 18 (the rest) independent experiments. Red asterisks indicate hits. More information is in Table S2. Please note that 5 compounds were not shown since 4 compounds were toxic and 1 was not available to retest. (C) Summary of the hit discovery rate of this study. See also Figure 6.

**Figure S6. Benidipine Rescues the Miro1 Phenotype.** (A-B) Similar to Figure S5, Miro1 intensity is measured in conditions indicated. (A) n=4 independent experiments. (B) n=3 independent experiments. (C) Fibroblasts were treated as indicated and run in SDS-PAGE. Quantification of the band intensity to that of GAPDH from the same blot. Compared to the far-left bar. n=4 independent experiments. Further explanation: Here we depolarized mitochondria with a different uncoupler, CCCP [1], instead of FCCP, because it had been used successfully in Western blotting on Miro1. We detected Miro1 and mitochondrial markers at 6 and 14 hours after CCCP treatment. We had previously demonstrated that in healthy control fibroblasts following CCCP treatment, Miro1 was degraded earlier (6 hours) than multiple other mitochondrial markers (14 hours) [1,2], consistent with the observation of proteasomal degradation of Miro1 prior to mitophagy [1-4]. This figure confirmed that both Miro1 degradation and subsequent damaged mitochondrial clearance were impaired in the PD cell line we used for screens. Importantly, Benidipine promoted Miro1 degradation after 6 hours following CCCP treatment and facilitated mitochondrial clearance as was evidenced by the degradation of ATP5β at 14 hours post-treatment. (D) RT-qPCR results of *Miro1* normalized to *GAPDH* from fibroblasts treated as indicated. n=3. (E) ATP levels were measured in fibroblasts as indicated. n=5. (F) Right: Representative confocal images of TMRM staining in fibroblasts. Left: Quantification of the TMRM intensity normalized to background. Scale bar: 25 μm. n=3 independent coverslips. (A, C, D, E, F) One-Way Anova Post Hoc Tukey Test. (G) Neurons were treated as indicated and Miro1 protein was detected by an ELISA normalized to the total protein amount [5,6] (Figure S7C-D). n=4 for DFP treated, 6 for the rest. Two-tailed Welch's T Test. Comparison within each cell line with "untreated", except otherwise labeled. Note that at baseline, Miro1 protein amount is increased in PD neurons, as reported in [7]. The same volume of solvent was added to controls. See also Figure 7.

**Figure S7. Miro1 in PBMCs and Miro1's Relation with Selective T-Type $Ca^{2+}$-Channel Blockers.** (A) PBMCs from a healthy donor (SBC, Table S4) were cultured and treated as indicated, and lysates were run in SDS-PAGE. (B) From blots as in (A), the band intensity is quantified by normalizing it to GAPDH from the same blot. n=3 independent experiments for Miro1 and 4 for ATP5β and VDAC. Two-tailed T Test. Left: Antimycin A and Oligomycin. Right: CCCP. (C) A representative standard curve using the Miro1 ELISA kit. Sigmoidal 4PL is used. (D) Intraplate and interpolate variability of the ELISA kit was demonstrated to be low. Intraplate: the same fibroblast cell lysate was run 4 times in the same plate. Interplate: the same protein standard (1.25 ng/ml) was run in 4 different plates. (E-F) ROC plots for Miro1 Ratio alone (E) and Miro1 Ratio combined with UPDRS (F). (G) Similar to Figure S5, fibroblasts were treated as indicated and Miro1 and Dapi were imaged. Heat map shows the mean Miro1 protein intensities calculated from the images. Each mean Miro1 value is expressed as a percentage of the mean Miro1 value from the same cell line with DMSO treatment alone. n=3-10 independent experiments. Precise p and n are in the figure. Two-tailed Mann-Whitney Test, compared to "DMSO alone". (H) Neurons were transfected with siRNA and treated as indicated. Miro1 was detected by ELISA as above. Comparison with "untreated" within the same line unless labeled. n=4. Two-tailed Welch's T Test. Compared to non-targeting (NT) siRNA, remaining *Cav3.2* after *Cav3.2* siRNA was 37.5-39.0% by RT-qPCR. See also Figure 7.

**SUPPLEMENTARY TABLE LEGENDS**
**Table S1: Demographic Information of Postmortem Brain Donors.**
**Table S2: Hit Compounds from the HTP Screening.**
**Table S3: Pathway Analysis.**
**Table S4: Demographic Information of PBMC Donors and Miro1 Values.**
**Table S5: Human Genetic Analysis.**

#### References

1. Hsieh, C.H., Li, L., Vanhauwaert, R., Nguyen, K.T., Davis, M.D., Bu, G., Wszolek, Z.K., and Wang, X. (2019). Miro1 Marks Parkinson's Disease Subset and Miro1 Reducer Rescues Neuron Loss in Parkinson's Models. Cell metabolism *30*, 1131-1140 e1137. 10.1016/j.cmet.2019.08.023.
2. Hsieh, C.H., Shaltouki, A., Gonzalez, A.E., Bettencourt da Cruz, A., Burbulla, L.F., St Lawrence, E., Schule, B., Krainc, D., Palmer, T.D., and Wang, X. (2016). Functional Impairment in Miro Degradation and Mitophagy Is a Shared Feature in Familial and Sporadic Parkinson's Disease. Cell stem cell *19*, 709-724. 10.1016/j.stem.2016.08.002.
3. Chan, N.C., Salazar, A.M., Pham, A.H., Sweredoski, M.J., Kolawa, N.J., Graham, R.L., Hess, S., and Chan, D.C. (2011). Broad activation of the ubiquitin-proteasome system by Parkin is critical for mitophagy. Human molecular genetics *20*, 1726-1737. 10.1093/hmg/ddr048.
4. Wang, X., Winter, D., Ashrafi, G., Schlehe, J., Wong, Y.L., Selkoe, D., Rice, S., Steen, J., LaVoie, M.J., and Schwarz, T.L. (2011). PINK1 and Parkin target Miro for phosphorylation and degradation to arrest mitochondrial motility. Cell *147*, 893-906. 10.1016/j.cell.2011.10.018.
5. Nguyen, D., Bharat, V., Conradson, D.M., Nandakishore, P., and Wang, X. (2021). Miro1 Impairment in a Parkinson's At-Risk Cohort. Front Mol Neurosci *14*, 734273. 10.3389/fnmol.2021.734273.
6. Bharat, V., Hsieh, C.H., and Wang, X. (2021). Mitochondrial Defects in Fibroblasts of Pathogenic MAPT Patients. Front Cell Dev Biol *9*, 765408. 10.3389/fcell.2021.765408.

7. Shaltouki, A., Hsieh, C.H., Kim, M.J., and Wang, X. (2018). Alpha-synuclein delays mitophagy and targeting Miro rescues neuron loss in Parkinson's models. Acta Neuropathol *136*, 607-620. 10.1007/s00401-018-1873-4.

### Figure S1

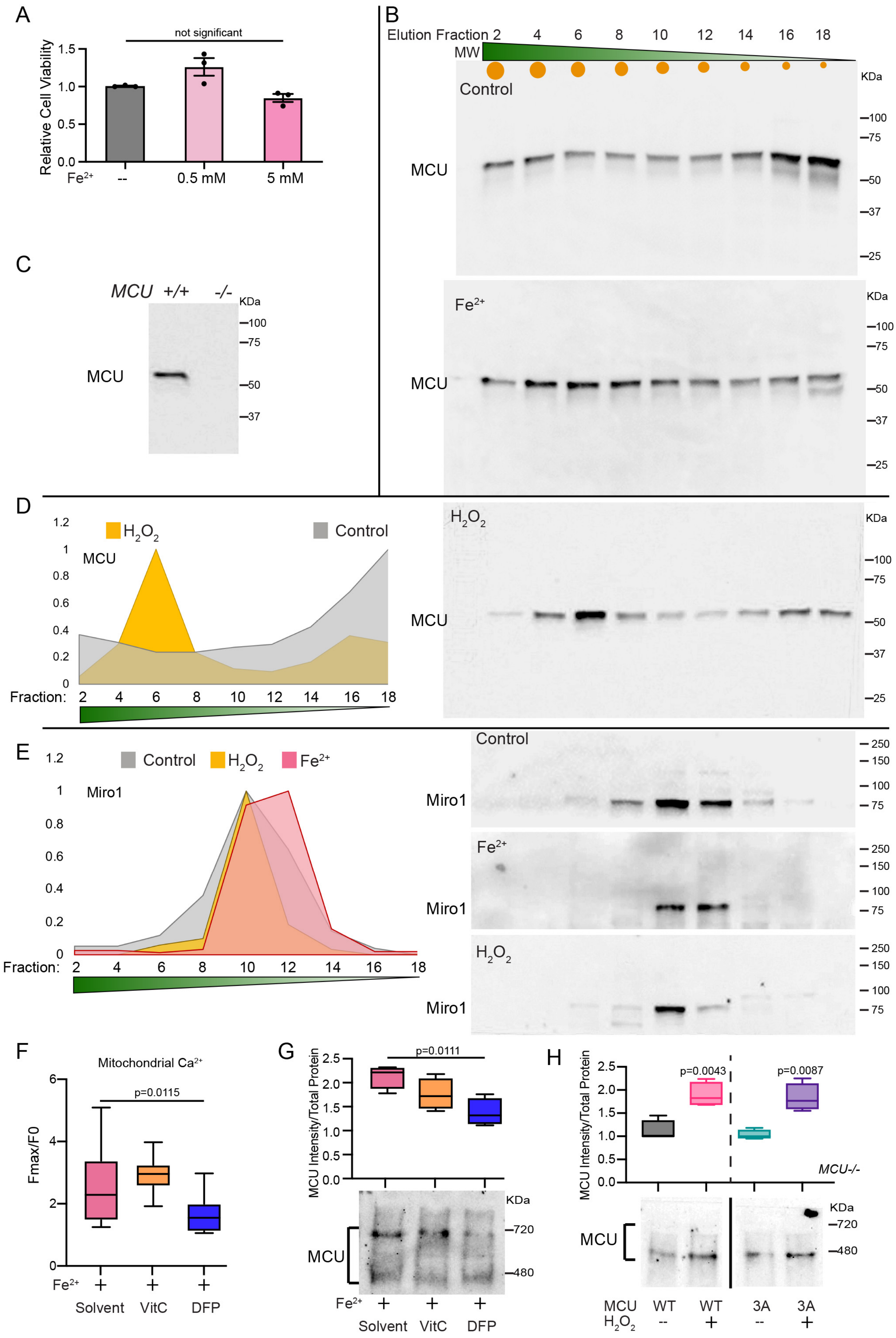

A
not significant
Relative Cell Viability
1.5
1.0
0.5
0.0
Fe2+
--
0.5 mM
5 mM
B
Elution Fraction 2 4 6 8 10 12 14 16 18
MW
Control
KDa
100
75
MCU
50
37
25
Fe2+
KDa
100
75
MCU
50
37
25
C
MCU +/+ -/-
KDa
100
75
MCU
50
37
D
H2O2
Control
MCU
1.2
1
0.8
0.6
0.4
0.2
0
Fraction: 2 4 6 8 10 12 14 16 18
H2O2
KDa
100
75
MCU
50
37
25
E
Control
H2O2
Fe2+
Miro1
1.2
1
0.8
0.6
0.4
0.2
0
Fraction: 2 4 6 8 10 12 14 16 18
Control
Miro1
250
150
100
75
Fe2+
Miro1
H2O2
Miro1
F
Mitochondrial Ca2+
p=0.0115
Fmax/F0
6
4
2
0
Fe2+
Solvent
VitC
DFP
G
p=0.0111
MCU Intensity/Total Protein
2.5
2.0
1.5
1.0
0.5
0.0
KDa
720
480
MCU
Fe2+
Solvent
VitC
DFP
H
p=0.0043
p=0.0087
MCU Intensity/Total Protein
MCU-/-
KDa
720
480
MCU
MCU WT WT 3A 3A
H2O2 -- + -- +

### Figure S2

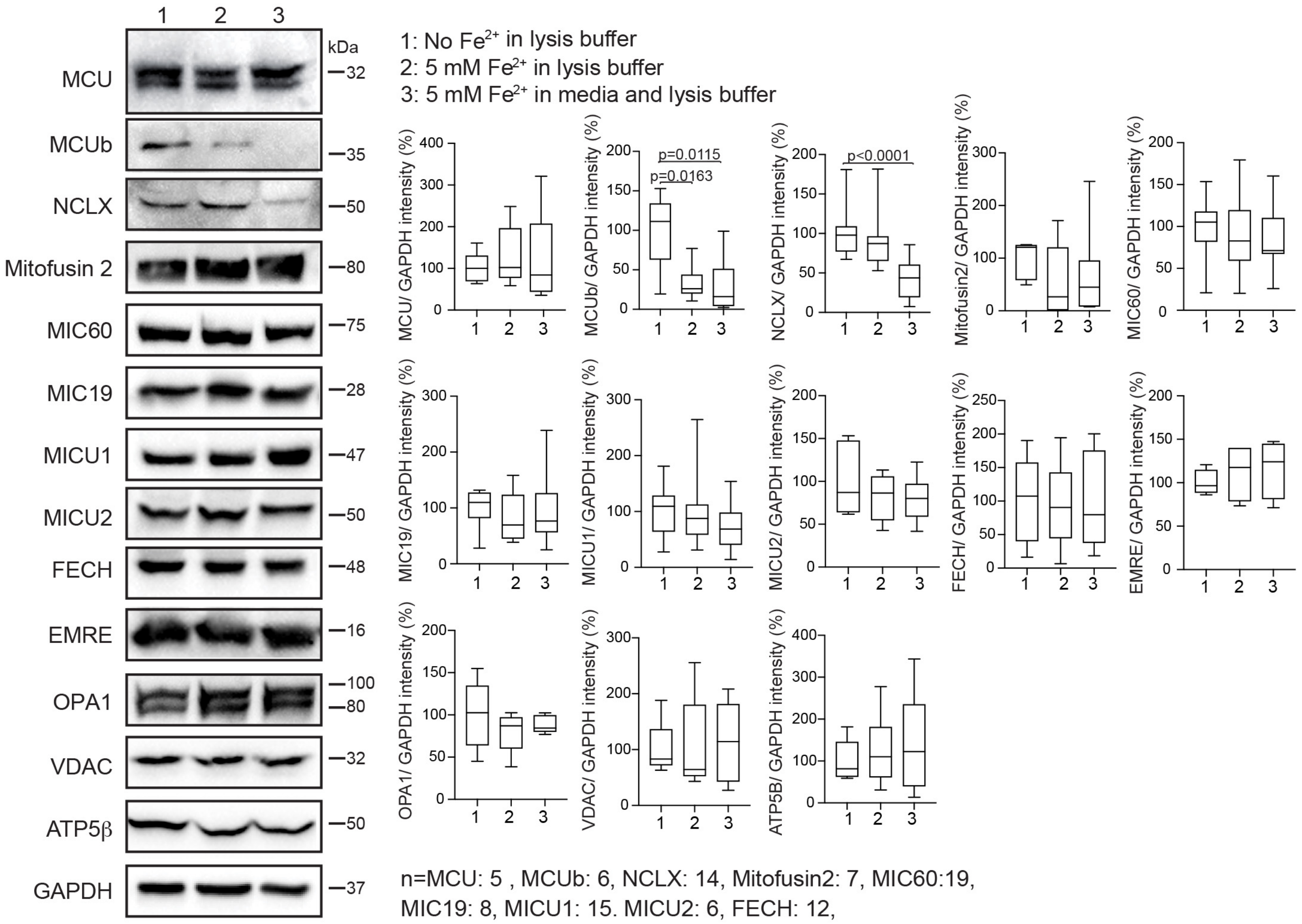


n=MCU: 5 , MCUb: 6, NCLX: 14, Mitofusin2: 7, MIC60:19, MIC19: 8, MICU1: 15. MICU2: 6, FECH: 12, EMRE: 4, OPA1: 5, VDAC: 5, ATP5β: 6.

### Figure S3

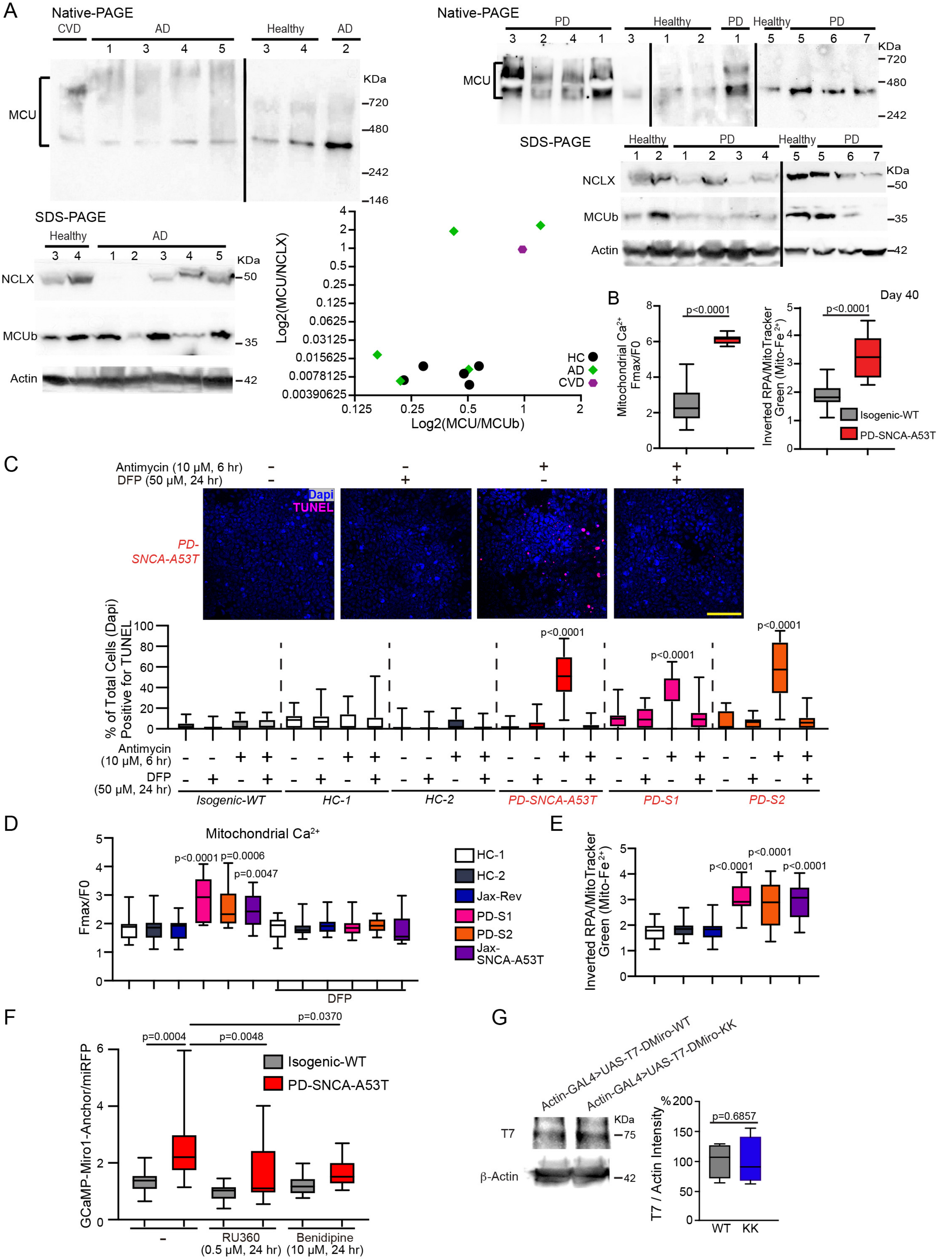

A
Native-PAGE
CVD
AD
Healthy
MCU
KDa
720
480
242
146
PD
Healthy
SDS-PAGE
NCLX
MCUb
Actin
50
35
42
Log2(MCU/NCLX)
Log2(MCU/MCUb)
HC
AD
CVD
B
Day 40
Mitochondrial Ca2+ Fmax/F0
Inverted RPA/MitoTracker Green (Mito-Fe2+)
p<0.0001
Isogenic-WT
PD-SNCA-A53T
C
Antimycin (10 μM, 6 hr)
DFP (50 μM, 24 hr)
Dapi
TUNEL
PD-SNCA-A53T
% of Total Cells (Dapi) Positive for TUNEL
Isogenic-WT
HC-1
HC-2
PD-SNCA-A53T
PD-S1
PD-S2
D
Mitochondrial Ca2+
Fmax/F0
p<0.0001
p=0.0006
p=0.0047
DFP
HC-1
HC-2
Jax-Rev
PD-S1
PD-S2
Jax-SNCA-A53T
E
Inverted RPA/MitoTracker Green (Mito-Fe2+)
F
GCaMP-Miro1-Anchor/miRFP
p=0.0004
p=0.0048
p=0.0370
Isogenic-WT
PD-SNCA-A53T
RU360 (0.5 μM, 24 hr)
Benidipine (10 μM, 24 hr)
G
Actin-GAL4>UAS-T7-DMiro-WT
Actin-GAL4>UAS-T7-DMiro-KK
T7
β-Actin
KDa
75
42
T7 / Actin Intensity %
p=0.6857
WT
KK

### Figure S4

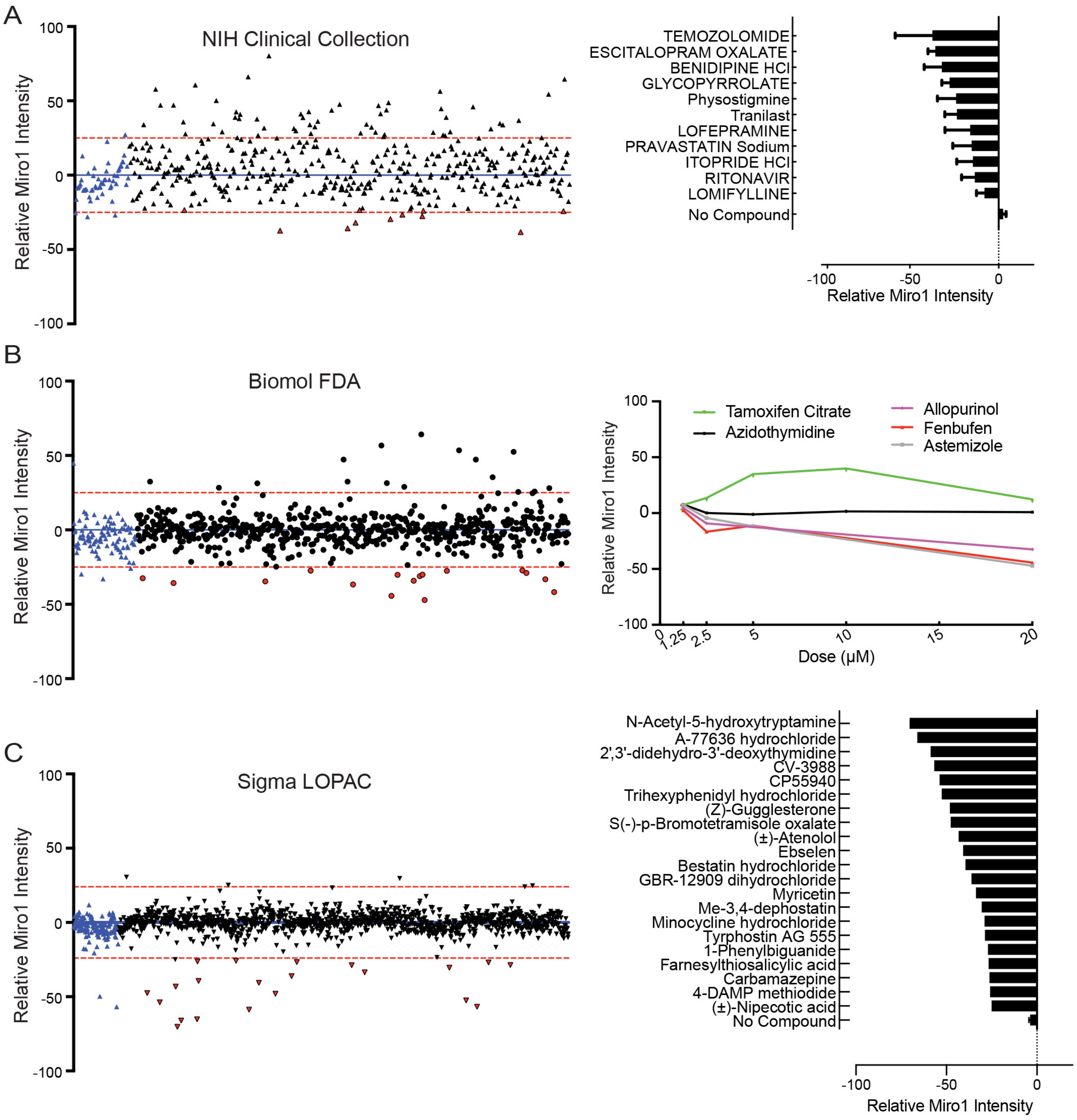
A
NIH Clinical Collection
Relative Miro1 Intensity
TEMOZOLOMIDE
ESCITALOPRAM OXALATE
BENIDIPINE HCl
GLYCOPYRROLATE
Physostigmine
Tranilast
LOFEPRAMINE
PRAVASTATIN Sodium
ITOPRIDE HCl
RITONAVIR
LOMIFYLLINE
No Compound
B
Biomol FDA
Tamoxifen Citrate
Azidothymidine
Allopurinol
Fenbufen
Astemizole
Dose (µM)
C
Sigma LOPAC
N-Acetyl-5-hydroxytryptamine
A-77636 hydrochloride
2',3'-didehydro-3'-deoxythymidine
CV-3988
CP55940
Trihexyphenidyl hydrochloride
(Z)-Guggleslerone
S(-)-p-Bromotetramisole oxalate
(±)-Atenolol
Ebselen
Bestatin hydrochloride
GBR-12909 dihydrochloride
Myricetin
Me-3,4-dephostatin
Minocycline hydrochloride
Tyrphostin AG 555
1-Phenylbiguanide
Farnesylthiosalicylic acid
Carbamazepine
4-DAMP methiodide
(±)-Nipecotic acid
No Compound

### Figure S5

A

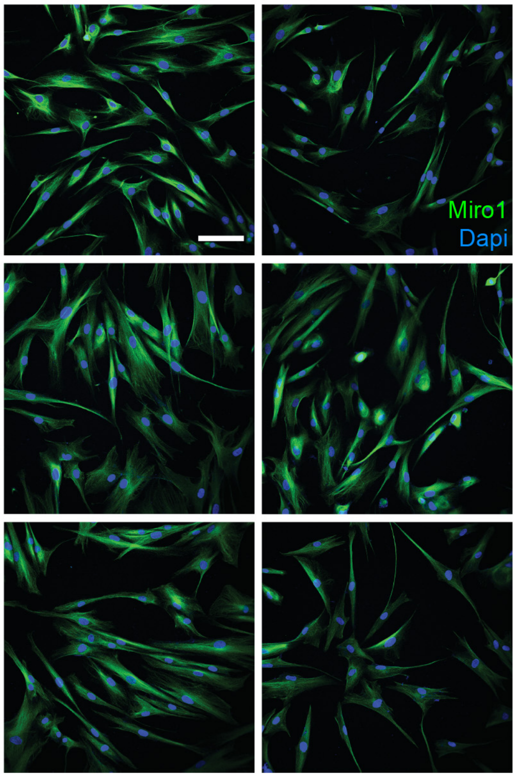


B

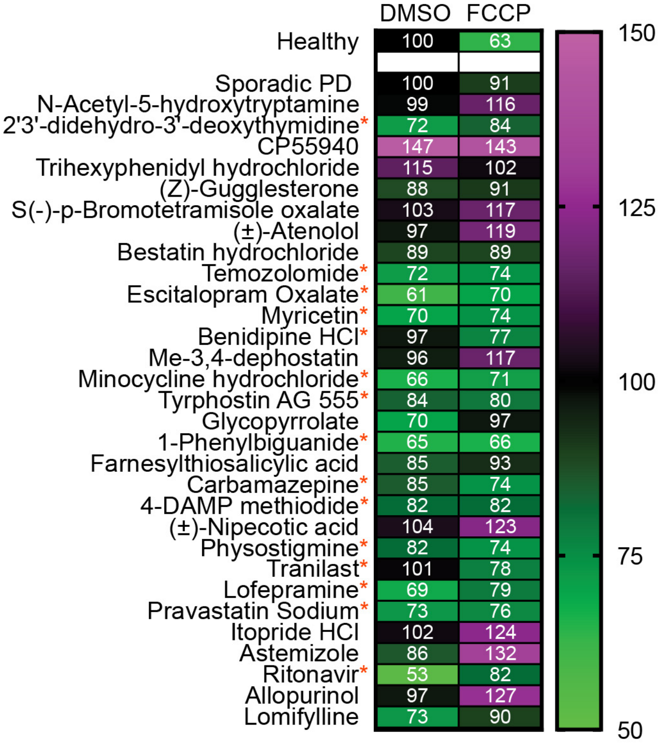


C

| **Library** | **Unique compounds** | **Screening concentration (μM)** | **Biological replicates** | **Primary Hits** | **Primary Hit rate (%)** | **Confirmed Hits** | **Confirmed Hit rate (%)** |
|---|---|---|---|---|---|---|---|
| NIH CC | 377 | 10 | 4 | 11 | 2.92 | 8 | 72.73 |
| Biomol FDA | 175 | 1.25-2.5-5-10-20 | 2 | 3 | 1.71 | 0 | 0 |
| Sigma LOPAC | 1269 | 20 | 1 | 21 | 1.65 | 7 | 33.33 |
| **Total** | 1821 | | | 35 | 1.92 | 15 | 42.86 |

*fresh compounds of only 34 out of 35 hits were available and tested

### Figure S6

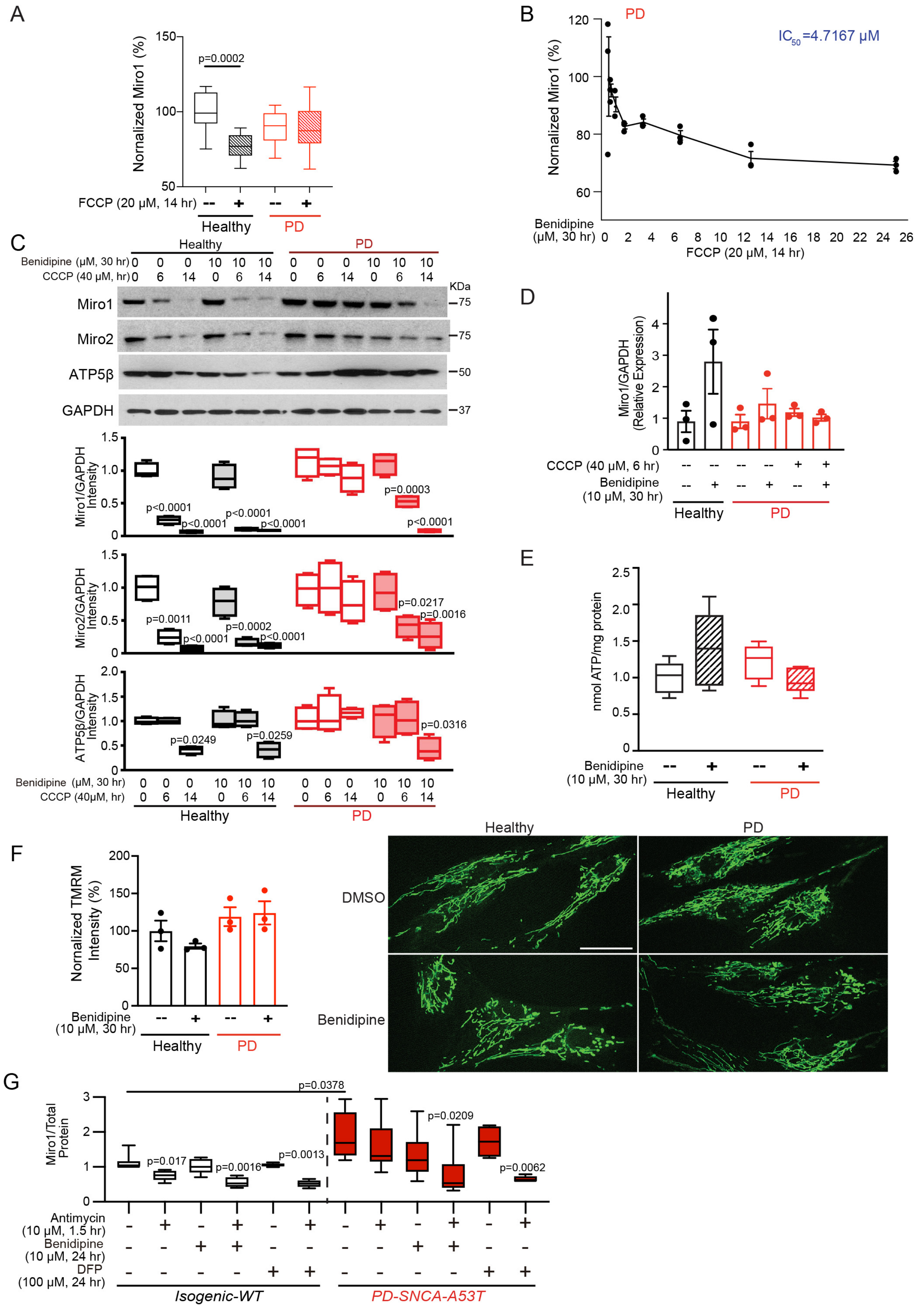

A
Nornalized Miro1 (%)
p=0.0002
FCCP (20 μM, 14 hr)
Healthy
PD
B
PD
IC50=4.7167 μM
Normalized Miro1 (%)
Benidipine (μM, 30 hr)
FCCP (20 μM, 14 hr)
C
Healthy
PD
Benidipine (μM, 30 hr)
CCCP (40 μM, hr)
KDa
Miro1
Miro2
ATP5β
GAPDH
Miro1/GAPDH Intensity
Miro2/GAPDH Intensity
ATP5β/GAPDH Intensity
D
Miro1/GAPDH (Relative Expression)
CCCP (40 μM, 6 hr)
Benidipine (10 μM, 30 hr)
Healthy
PD
E
nmol ATP/mg protein
Benidipine (10 μM, 30 hr)
Healthy
PD
F
Nornalized TMRM Intensity (%)
Benidipine (10 μM, 30 hr)
Healthy
PD
DMSO
Benidipine
G
Miro1/Total Protein
p=0.0378
Antimycin (10 μM, 1.5 hr)
Benidipine (10 μM, 24 hr)
DFP (100 μM, 24 hr)
Isogenic-WT
PD-SNCA-A53T

### Figure S7

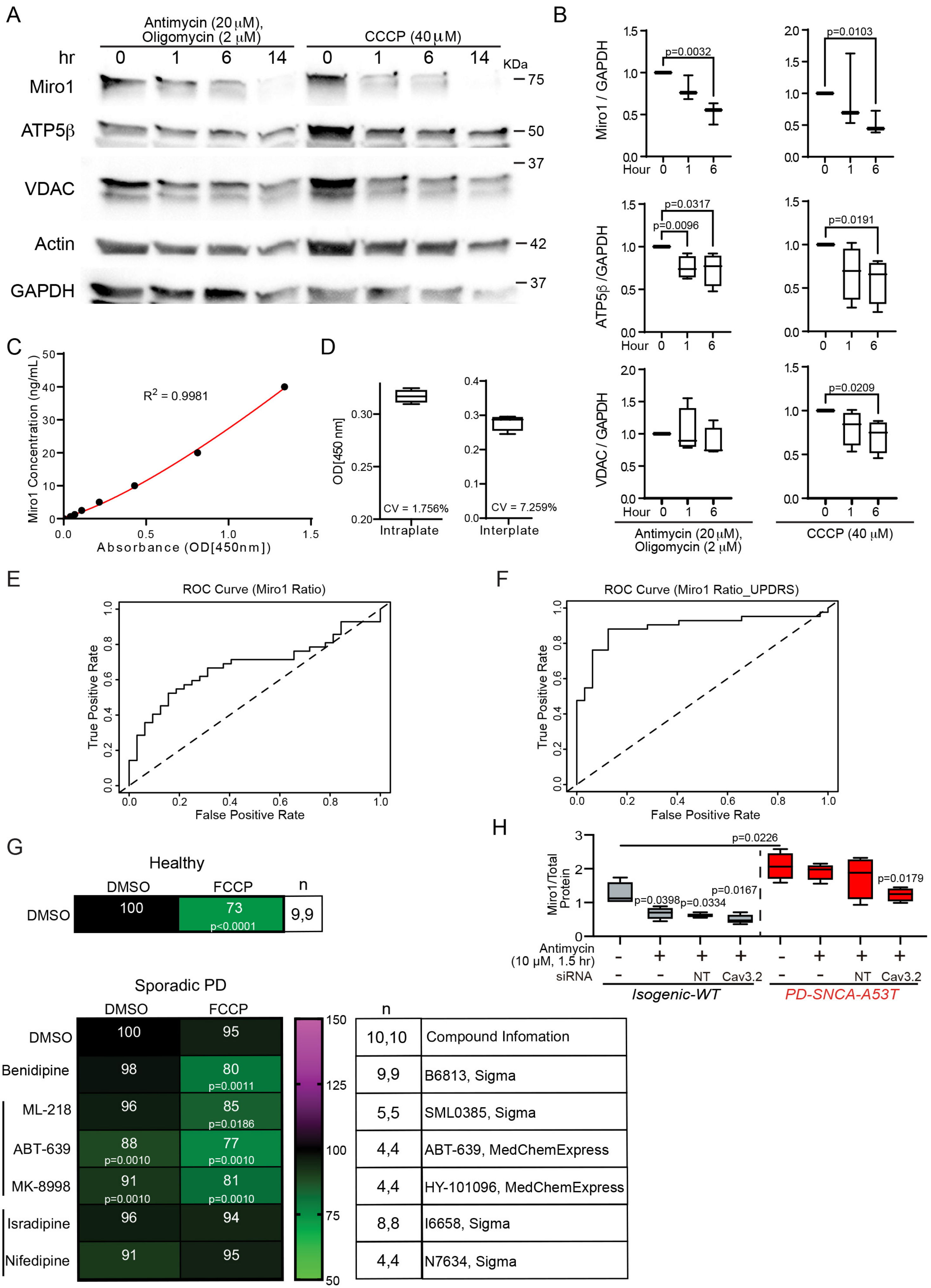
A
Antimycin (20 μM), Oligomycin (2 μM)
CCCP (40 μM)
hr 0 1 6 14 0 1 6 14
KDa
Miro1 — 75
ATP5β — 50
— 37
VDAC
Actin — 42
GAPDH — 37
B
Miro1 / GAPDH
p=0.0032
p=0.0103
ATP5β /GAPDH
p=0.0317
p=0.0096
p=0.0191
VDAC / GAPDH
p=0.0209
Hour 0 1 6
Antimycin (20 μM), Oligomycin (2 μM)
CCCP (40 μM)
C
Miro1 Concentration (ng/mL)
$R^2$ = 0.9981
Absorbance (OD[450nm])
D
OD[450 nm]
CV = 1.756%
Intraplate
CV = 7.259%
Interplate
E
ROC Curve (Miro1 Ratio)
True Positive Rate
False Positive Rate
F
ROC Curve (Miro1 Ratio_UPDRS)
True Positive Rate
False Positive Rate
G
Healthy
DMSO FCCP n
DMSO 100 73 p<0.0001 9,9
Sporadic PD
DMSO FCCP
DMSO 100 95
Benidipine 98 80 p=0.0011
T ML-218 96 85 p=0.0186
ABT-639 88 p=0.0010 77 p=0.0010
MK-8998 91 p=0.0010 81 p=0.0010
L Isradipine 96 94
Nifedipine 91 95
150 125 100 75 50
n
10,10 Compound Infomation
9,9 B6813, Sigma
5,5 SML0385, Sigma
4,4 ABT-639, MedChemExpress
4,4 HY-101096, MedChemExpress
8,8 I6658, Sigma
4,4 N7634, Sigma
H
Miro1/Total Protein
p=0.0226
p=0.0398
p=0.0334
p=0.0167
p=0.0179
Antimycin (10 μM, 1.5 hr)
siRNA
NT Cav3.2
Isogenic-WT
PD-SNCA-A53T
